# Evolved bacterial resistance to the chemotherapy gemcitabine modulates its efficacy

**DOI:** 10.1101/2022.09.07.506952

**Authors:** Serkan Sayin, Brittany Rosener, Carmen G Li, Bao Ho, Olga Ponomarova, Doyle V Ward, Albertha JM Walhout, Amir Mitchell

## Abstract

Drug metabolism by the microbiome can influence anti-cancer treatment success. We previously suggested that chemotherapies with antimicrobial activity can select for adaptations in bacterial drug metabolism that can inadvertently influence the host’s chemoresistance. We demonstrated that evolved resistance against fluoropyrimidine chemotherapy lowered its efficacy in worms feeding on drug-evolved bacteria (Rosener et al., 2020). Here we examine a model system that captures local interactions that can occur in the tumor microenvironment. Gammaproteobacteria colonizing pancreatic tumors can degrade the nucleoside-analog chemotherapy gemcitabine and, in doing so, can increase the tumor’s chemoresistance. Using a genetic screen in *Escherichia coli*, we mapped all loss-of-function mutations conferring gemcitabine resistance. Surprisingly, we found that one third of resistance mutations increase or decrease bacterial drug breakdown and therefore can either lower or raise the gemcitabine load in the local environment. Experiments in three *E. coli* strains revealed that evolved adaptation converged to inactivation of the nucleoside permease NupC, an adaptation that increased the drug burden on co-cultured cancer cells. The two studies provide complementary insights on the potential impact of microbiome adaptation to chemotherapy by showing that bacteria-drug interactions transpire locally and systemically and can influence chemoresistance in the host.

## Introduction

Clinical research on the influence of intratumor bacterial infection can be dated back to more than 150 years ago^1^. However, in the past decade, research of the tumor-microbiome gained significant momentum with the maturation of DNA sequencing technologies and advancement of microbiome research. Multiple resent studies of bacterial colonization in human tumors outlined the magnitude of this phenomenon (reviewed here^1–3^) Collectively, these works establish that the proportion of infected tumors greatly varies across tumor types and that many tumors harbor microbiomes with a distinctive and characteristic composition of bacterial species. In some cases, as in breast and pancreatic cancer, more than 60% of tumors harbored a tumor-microbiome^4^. The microbiome, in turn, is known to influence cancer disease through multiple independent mechanisms, including promotion of neoplastic processes in healthy host cells, modulation of the host anti-tumor immune response, and by bacterial biotransformation of anticancer drugs^3,5–7^

Bacterial metabolism of xenobiotics, including breakdown of host-targeted drugs, is prevalent^8^. Estimates from recent drug screens show that two thirds of human-targeted drugs can be metabolized by at least one bacterial species that is present in the human gut microbiome^9^. Yet, these interactions are reciprocal, bacteria both metabolize the host-targeting drugs and are also frequently impacted by them^10^. Roughly 25% of host-targeted drugs are potent inhibitors of bacterial growth at physiological concentrations^11^. This proportion is doubled for antineoplastic drugs and almost all anticancer drugs that belong to the antimetabolite drug class have potent antimicrobial activity^11^. A key underexplored question that arises from these reciprocal drug-microbiome interactions is how they impact one another given the ability of microorganisms to evolve and change over short time scales within the host^12–16^. Specifically, given that bacteria often rapidly evolve resistance to antimicrobial drugs, it is plausible that adaptation to host-targeting drugs that are also antimicrobial will alter bacterial drug metabolism or its transport^17,18^. Such adaptions have been repeatedly observed with standard antibiotics^19^. In such cases, evolved resistance in tumor-colonizing bacteria may increase or decrease drug availability to the tumor cells which, in turn, may interfere with the efficacy of the chemotherapy.

The tumor-microbiome in pancreatic cancer has attracted much attention recently due to the prevalence of infection in pancreatic ductal adenocarcinoma (PDAC)^4,7,20–23^. Studies have uncovered multiple independent mechanisms through which microbes influence oncogenesis^22,23^, disease progression^7^ and treatment success^21^ in the pancreas. Bacterial infection is attributed to retrograde bacterial migration from the gastrointestinal tract into the pancreas^20,23^. Characterization of the PDAC tumor-microbiome by 16S rRNA gene sequencing showed that proteobacteria are highly enriched relative to the gut microbiome and that they are highly prevalent in pancreatic tumors^4,21,23^. Recent work suggested that pancreatic colonization can impede therapy with gemcitabine (2’, 2’-difluoro 2’ deoxycytidine, dFdC), a front-line chemotherapy drug that is used for PDAC treatment^21^. Further clinical data provided circumstantial evidence indicating that this interaction may indeed take place in treated patients^24–26^.

Gemcitabine drug metabolism is well-understood in the model gamma-proteobacteria *E. coli*^27^ (Figure 1A). The antimetabolite gemcitabine, a nucleoside analog, is imported into the bacterial cell through the nucleoside transporter NupC and is then phosphorylated. Gemcitabine triphosphate can be incorporated into a newly synthesized DNA strand and then can interfere with chain elongation by masked chain termination (similar to mammalian cells^28^). Therefore, despite its clinical use as an anticancer drug, gemcitabine’s mechanism of action potentially makes it a broadly toxic, antimicrobial compound. Previous works showed that some bacterial species can rapidly convert gemcitabine into the less toxic metabolite 2’,2’-difluoro-2’-deoxyuridine (dFdU)^21,29,30^. In gamma-proteobacteria, gemcitabine degradation proceeds through a specific isoform of the cytidine deaminase enzyme (Cdd_L_)^21^. The well-characterized interactions between tumor cells, gemcitabine and gamma-proteobacteria puts forth a good model system for testing how bacterial adaptation can impact drug metabolism and potentially influence the tumor’s chemoresistance.

**Figure 1:**
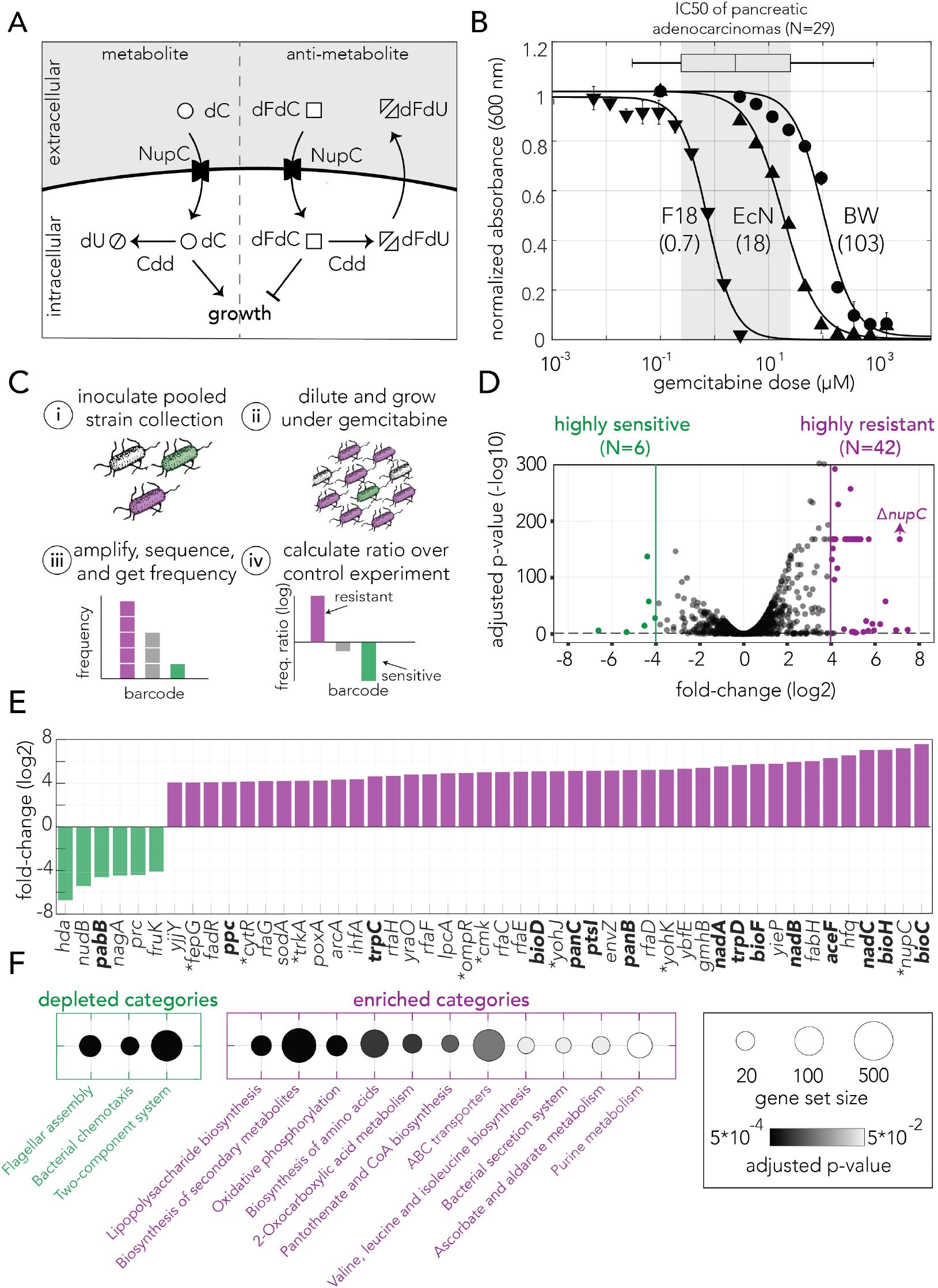
Genetic screen identifies gemcitabine sensitive and resistant loss-of-function mutations in *E. coli*. **A**. Gemcitabine transport and metabolism in *E. coli*. Similar to deoxycytidine (dC), gemcitabine (dFdC) is imported into the cell through the nucleoside permease NupC. Intracellular gemcitabine is either converted into the less toxic metabolite dFdU the cytidine deaminase Cdd or is phosphorylated and incorporated into the DNA. **B**. Gemcitabine dose response curves for three *E. coli* strains (three technical replicates, performed once). The inferred IC50 are shown in parenthesis. Gemcitabine IC50 range of 29 pancreatic adenocarcinomas are shown as a box plot above the graph. **C**. Overview of the pooled screening approach. Pooled cultures of the knockout strain collection were inoculated (i) and grown for multiple hours with or without gemcitabine (ii), DNA extracted from cells was used to amplify, sequence and calculate the frequency of barcodes that correspond to individual strains (iii), The ratio of barcode frequencies of each strain in gemcitabine and control conditions were used to identify sensitive and resistant knockouts (iv). Experiment includes three biological replicates (independent inoculums) and performed once. **D**. Volcano plot of the genetic screen results.Green and purple dots represent sensitive and resistant knockout strains, respectively. **E**. Highly sensitive and resistant strains. Asterisk sign marks the gene knockouts likely involved in gemcitabine transport and phosphorylation. Knockouts marked in bold were characterized as slow growing strains. **F**. Statistically significant enriched and depleted KEGG categories identified by the genetic screen. The marker size shows the number of genes in the category and the gray scale marks the statistical significance.

Through a pooled genetic screen, we systematically mapped all loss-of-function mutations that increase *E. coli*’s resistance to gemcitabine and found that inactivation of over than forty genes increased bacterial resistance by over than 16-fold. This observation led us to conclude that resistance can rapidly emerge under natural selection through gene inactivation within a single evolutionary step. Using a functional assay, we found that one third of resistance mutations impacted extracellular drug concentrations. Co-culturing bacteria harboring these loss-of-function mutations with cancer cells confirmed that these adaptive mutations have the potential to alter chemoresistance of neighboring tumor cells. Finally, through in-vitro evolution we studied which adaptations emerge under drug selection in three *E. coli* strains. We found that inactivation of the drug transporter NupC arises in all evolved strains. This inactivation leads to decreased bacterial drug import and therefore reduces the rate of gemcitabine breakdown. Reduced bacterial breakdown, in turn, increases gemcitabine availability for neighboring tumor cells. Our work reveals that bacterial adaptation to the frontline chemotherapy drug gemcitabine can take place rapidly and can ultimately increase the chemosensitivity of the hosting tumor. Given that administration of antibiotics can be detrimental to cancer patients^24,26,31,32^, our work is potentially impactful since highlights that tumor colonization by chemotherapy metabolizing bacteria may not hamper anticancer treatment in the long run.

## Results

### The *E. coli* Resistome against gemcitabine

We first set out to determine the inhibitory concentration of gemcitabine in three *E. coli* strains: BW25113 (a K-12 lab strain), F-18 (a human fecal isolate), and Nissle 1917 (a human fecal isolate that is used as probiotic). We characterized the inhibitory concentrations by monitoring bacterial growth inhibition after 12 hours. Figure 1B shows the sensitivity curves and the inhibitory concentration that reduced culture density by 50% (IC50). We observed that gemcitabine can completely inhibit the growth of all three strains, with F-18 being most sensitive (IC50=0.7 μM) and BW25113 being most resistant (IC50=103 μM). These concentrations are comparable to the IC50 reported for 29 pancreatic adenocarcinomas in the GDSC2 dataset^33^ (boxplot above the graph in figure 1B).

We next conducted a genetic screen with a collection of 3,680 single-gene knockout strains to systematically map all non-essential genes that influence gemcitabine resistance when depleted (typically referred to as the drug resistome). We used a pooled screening approach that we recently developed which relies on sequencing DNA barcodes that are unique for each gene-knockout in the strain collection^17,34^. Figure 1C outlines the main steps of the screening method: the pooled knockout strains were inoculated and grown in media containing a high gemcitabine concentration (140 μM) while a control culture was inoculated in media without drug. Once the cultures reached a late logarithmic growth phase, cells were lysed, and DNA was extracted for amplification and sequencing of the barcode region. Lastly, we compared the frequency of each barcode, corresponding to an individual gene-knockout, in the drug and control conditions. This comparison revealed strains whose frequency was significantly increased or decreased in gemcitabine. Such enrichment or depletion correspond to increased or decreased drug resistance, respectively. We used three biological replicates to infer statistical significance. We identified over a million and a half barcode containing reads in each replicate that corresponded to roughly 3,500 unique barcodes (knockout strains). Supplementary Figure 1 shows detailed information on the screen coverage and replication quality.

Figure 1D shows a volcano plot representation of the screen results. Using very conservative cutoff values for fold-change (>16) and FDR-adjusted p-value (<0.05), we identified 42 gemcitabine-resistant knockout strains and 6 gemcitabine-sensitive strains (shown as purple and green circles in Figure 1D). The screen results appear in Supplementary table 1. Reassuringly, we found that knockout of the known drug transporter (*nupC*) is among the top resistors. When we inspected the gene annotation of the top resistors, we identified multiple hits from the known target pathway of the drug (Figure 1E). These included the permease (*nupC*), the transcriptional regulator *cytR* (a repressor of both *nupC* and *cdd*), and the cytidylate kinase (*cmk*) that likely phosphorylate intracellular gemcitabine. We also found multiple hits that encode membrane proteins or transporters, including *ompR* and *envZ* that together regulate permeability channels for nutrients, toxins, and antibiotics^35^. Lastly, we observed a high prevalence of genes coding for metabolic enzymes that can considerably slow down growth when mutated^34^. Indeed, a statistical test revealed that gene knockouts that were previously identified as reducing growth^36^ were highly enriched in the set of gemcitabine resistant strains (*p-value*= 8.6186e-18, Fisher exact test). The overlap between resistant strains and slow growing strains is sensible given that slow growth reduces the rate of DNA synthesis. Since gemcitabine is quickly degraded by the bacterial Cdd enzyme, it was only transiently present in the extracellular media during the screen experiment. Under such a transitory stress, slower consumption of the antimetabolite is likely beneficial (normal growers incorporated much more gemcitabine into their DNA and remain arrested while slow growers incorporated less and therefore avoid arrest).

Next, we used the gene set enrichment analysis tool GAGE^37^ to test for functional enrichment using the Kyoto Encyclopedia of Genes and Genomes (KEGG)^38^ and Gene Ontology (GO)^39^ databases. This analysis is complementary to the previous analysis, since it considers the enrichment values from all strains, rather than only the limited set of hit strains obtained by imposing strict cutoffs. Figure 1F shows the functional enrichment by KEGG pathways (enrichment indicates that pathway inactivation increased resistance). Reassuringly, we observed a high agreement between the functional enrichment by GAGE and the annotation of the top hits. Specifically, we observed enrichment in purine synthesis and membrane transporters as well as multiple metabolic pathways impacting bacterial growth rate^34^. Supplementary table 2 provides the full list of enriched and depleted categories.

Taken together the results from our genetic screen outline three adaptation strategies that increase bacteria’s gemcitabine resistance: reduced drug import by inactivating membrane proteins and transporter systems, changes in the drug metabolism through mutations in the target pathway, and inactivation of metabolic genes that slowdown bacterial growth. Importantly, since these resistance adaptations arise from knockout of single genes, they are all accessible within a single evolutionary step (e.g., a single gene inactivating mutation).

### The impact of bacterial resistance on bacterial drug degradation

Our genetic screen revealed alternative adaptation strategies that increase bacterial resistance against gemcitabine. However, for most loss-of-function mutations, it remains to be determined how they will influence the rate of bacterial drug degradation (and ultimately drug availability for neighboring cancer cells). We therefore designed a functional assay to detect changes in drug degradation rate relative to the wild-type strain (outlined in Figure 2A). We incubated a knockout strain of interest in saline with a high gemcitabine concentration and collected the conditioned supernatant after a short incubation period (15 or 45 minutes). We diluted the conditioned supernatant into regular media and monitored the growth of a drug-sensitive reporter strain (a *cdd* knockout that cannot degrade gemcitabine) in this media. Finally, we used growth curves of the reporter strain as a proxy for the gemcitabine concentration in the conditioned supernatant. We reasoned that a conditioned supernatant containing high drug concentration is indicative of slow degradation by the strain of interest. This difference in drug concentration will in turn manifest as slow growth of the reporter strain (blue curve in figure 2A). We note that this detection method is insufficient to resolve the mechanism underlying the slowed degradation since it only measures drug availability in the extracellular environment after incubation (e.g., both slow import and slow deamination will be interpreted as slow degradation).

**Figure 2:**
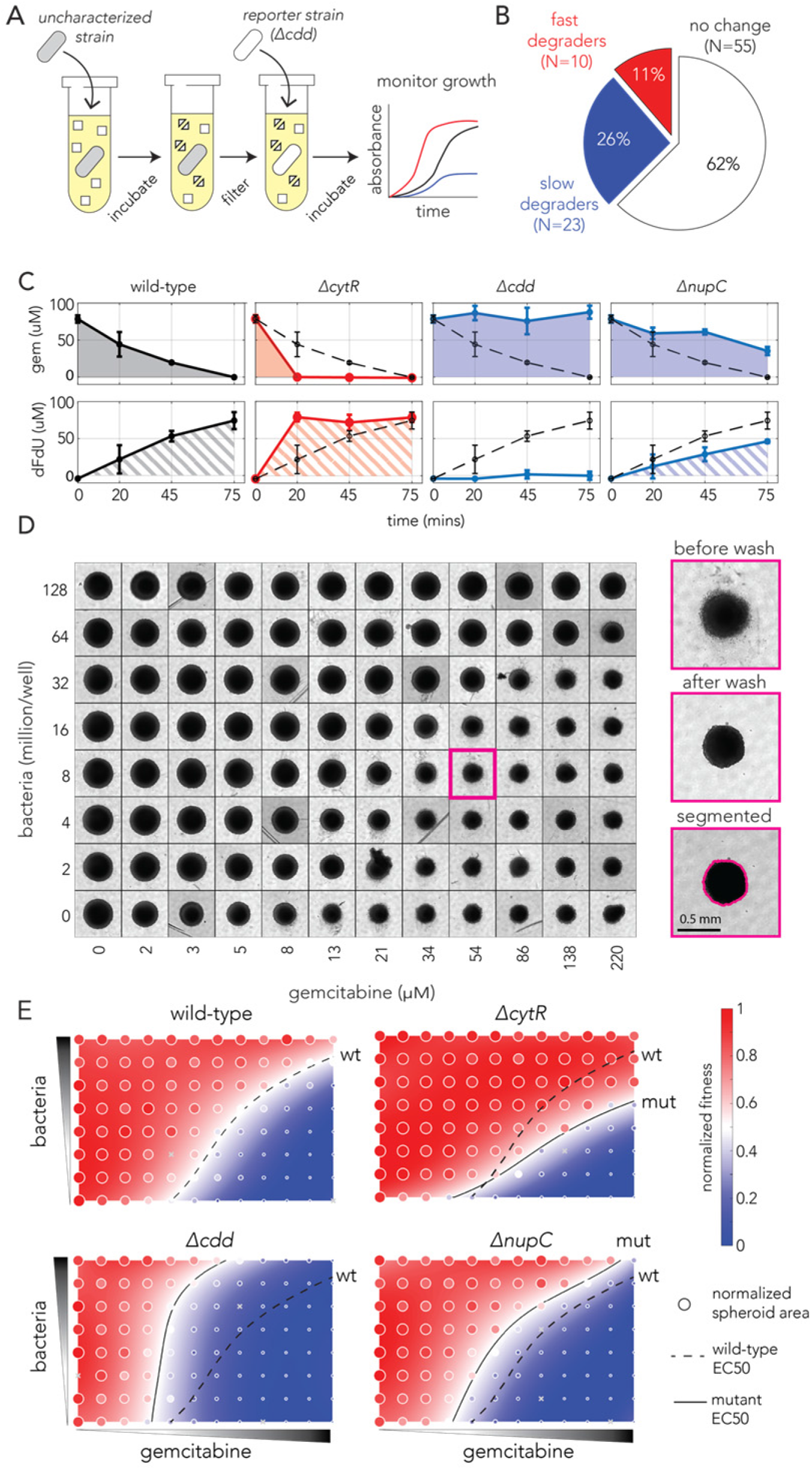
Bacterial gemcitabine resistance can oppositely affect drug degradation and impact neighboring cancer cells. **A**. The functional drug breakdown assay. Each knockout strain was inoculated in saline containing gemcitabine and incubated for 15 or 45 minutes. Conditioned supernatant was then filtered and mixed with fresh media before inoculating a reporter strain. Reporter strain growth was used as a proxy for changes in gemcitabine availability in the conditioned supernatant. **B**. Results of the drug breakdown assay for the top 88 resistant knockouts. A third of all tested knockouts (33/88) influenced the drug degradation rate. **C**. Chemical validation of the functional assay for the slowest and fastest degraders. GC-MS was used to measure gemcitabine concentration (shaded area, top panels) and its degradation product dFdU (hatched area, lower panels) in conditioned supernatant (colors as in A). The dashed black curve marks the measurements after incubation with the wild-type strain. The error bars show the standard deviation of three biological replicates. **D**. Co-culture experiments shows bacterial mutations can influence gemcitabine efficacy in neighboring spheroids of cancer cells. Representative microscopy images of spheroids that were co-cultured with wild-type bacteria across multiple concentrations of drug and bacteria (large images show the same spheroid before the wash, after the wash that removed dead cell debris, and after image segmentation. **E**. Bacterial mutations that modulate drug degradation can impact drug efficacy neighboring cancer cells. Results of spheroid experiments with the slowest and fastest degraders and the inferred fitness landscapes. Each panel shows the fitness landscape calculated from spheroid size across a range of gemcitabine and bacteria concentrations (as in D). The color code shows the normalized spheroid size (ranging from the smallest spheroid to the largest one). The dashed black line marks the parameter combination (bacteria and gemcitabine concentrations) that reduce spheroid growth by 50% (EC50 front) and solid lines show the EC50 front for the knockout strains. Shifts in the EC50 front, relative to the wild-type front, indicate that co-incubation with the knockout strain during drug exposure altered the spheroid’s chemo-resistance.

We evaluated the degradation rate of the top 42 resistors and additional 46 resistant strains (supernatant collection was repeated three times on different days). Figure 2B shows a summary of the results of this assay (supplementary Figure 2 shows observed growth curves). We found one third of the tested knockout strains modulated extracellular drug availability (33 of 88). Specifically, 11% of all strains were fast degraders and 26% of strains were slow degraders (one-tailed t-test, FDR-adjusted *p-value* < 0.1). We next decided to validate the conclusions of our functional assay using an independent chemical approach. We focused on the fastest and the slowest degraders. We incubated each strain with gemcitabine and sampled aliquots of the supernatant at predetermined timepoints (20, 45 and 75 minutes). We then used gas chromatography–mass spectrometry (GC-MS) to measure the concentration of gemcitabine and its degradation product in the conditioned supernatant (Figure 2C). In agreement with our functional assay, the GC-MS measurements confirmed that the gene knockouts indeed altered the availability of gemcitabine and in the extracellular environment. The increased availability of the drug breakdown product (dFdU) support the conclusion that rate of drug metabolism is underlying this change (as opposed to intracellular drug accumulation^40^)

Lastly, we tested if bacterial resistance mutations influence the drug sensitivity of co-cultured cancer cells. While previous works relied on sequential exposure to the drug^21,30^, using conditioned bacterial supernatant on cancer cells, we reasoned that a co-culture system will reveal if bacterial degradation is sufficient to impact drug efficacy in neighboring cancer cells that are simultaneously exposed to the drug. We co-cultured with spheroids of the CT-26 murine cancer cell-line and simultaneously treated them with gemcitabine for 4 hours (gemcitabine cytotoxicity can be recapitulated in this cell-line^21^). Spheroids were then washed and left to grow for a week in media supplemented with antibiotics. Lastly spheroids were washed again to remove dead cells and the area of the spheroids was measured to evaluate gemcitabine’s efficacy on the cancer cells. Figure 2D shows microscopy images of a representative microwell plate. As the figure shows we observed sensible trends in these experiments: First, we observed that spheroid size was inversely correlated with gemcitabine concertation (lower spheroid row in Figure 2D) and that bacterial concentration, without any drug, did not significantly impact spheroid growth (left spheroid column in Figure 2D). Reassuringly, we observed that co-cultured bacteria can mitigate gemcitabine damage and that the magnitude of rescue depended on the bacterial concentration (spheroid area is overall increased in upper spheroid rows in Figure 2D).

We used our systematic measurements of spheroid area to fit a fitness landscape. This procedure allowed us to infer the effective drug concentration that leads to a 50% change in area (EC50) for any bacterial concentration. Figure 2E shows the landscapes for the wild-type strain, a control non-degrader (*cdd* knockout) and the fastest and slowest degraders *(cytR* and *nupC knockouts*, respectively). As evidenced by the changes in the EC50 fronts, we observed that the spheroid fitness landscapes were considerably different depending on the co-cultured bacteria during drug exposure. In agreement with our functional and chemical assays, increased chemoresistance relative to the wild-type strain was observed when spheroids were co-incubated with the fast degrader (*cytR* knockout). In contrast, decreased chemoresistance relative to the wild-type strain was for co-incubation was with the slow degrader (*nupC* knockout). Additional experiments with nine more bacterial resistors showed they also impacted spheroid chemoresistance (Supplementary figure 3).

Taken together, the results of the spheroid experiments demonstrate that mutations impacting gemcitabine degradation rates in bacteria can indeed impact neighboring cancer cells simultaneously with the drug. Importantly, we observed that bacterial resistors can have opposite influences on gemcitabine sensitivity of co-cultured cancer cells. For examples, the *nupC* knockout decreased chemoresistance while the *cytR* knockout increased it. A key question remaining is which adaptations will naturally transpire during bacterial evolution under drug selection.

### Evolved bacterial resistance against gemcitabine

The genetic screen uncovered multiple loss-of-function mutations that confer bacterial gemcitabine resistance. Yet, such screens are insufficient for determining which gene inactivation, if any, will emerge under natural selection. Moreover, since evolution can leverage additional processes beyond gene-inactivation, such as gain-of-function, adaptation may follow an entirely different evolutionary trajectory. We applied the widely used serial transfer approach to select for evolved drug resistance in bacteria^41^. Such in-vitro experiments can shed light on the mechanisms underlying resistance and the time scale needed to acquire resistance. To test if reoccurring adaptations emerge, we used the three *E. coli* strains that were characterized by different drug sensitivity levels (Figure 1B). Once the serial transfer experiment ended, we evaluated if drug resistance increased in the population and isolated single resistant clones for whole genome-sequencing (Figure 3A).

**Figure 3:**
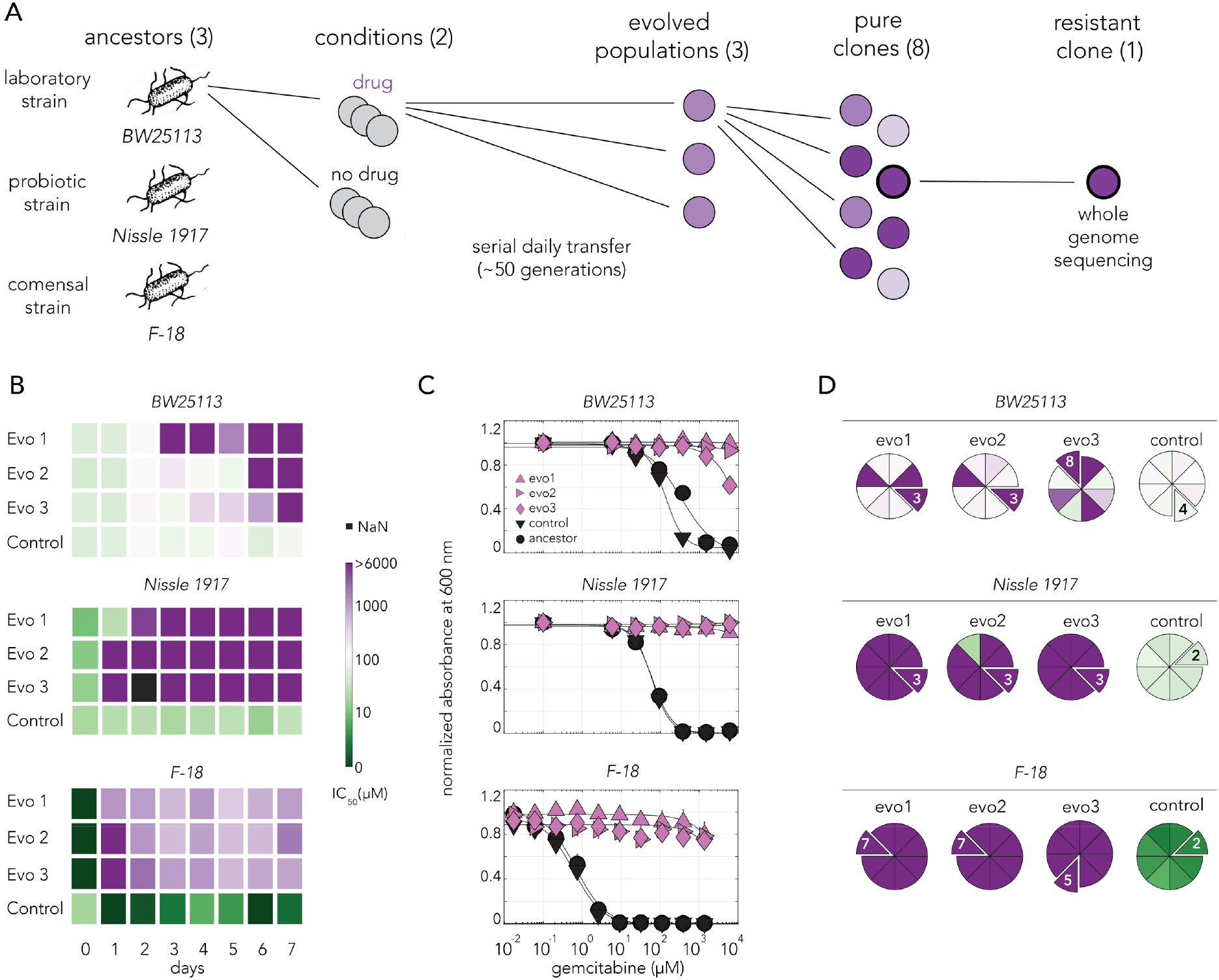
Gemcitabine selection leads to rapid evolved resistance in three *E. coli* strains. **A**. Overall approach for the lab evolution experiment. Three *E. coli* strains were evolved over 50 generations in serial transfer evolution experiment. We measured gemcitabine resistance of last day populations and eight single clones were isolated from each population. A single resistant clone was used for the whole genome sequencing and for identifying of the underlying mechanism for drug resistance. **B**. Heatmaps showing the temporal changes in gemcitabine IC50 of evolving populations in each day of the serial transfer experiment. **C**. Gemcitabine dose response curves of last day populations from lab evolution experiment. All of the gemcitabine evolved lines, but not the control evolved lines, developed a high resistance against gemcitabine. **D**. Pie charts showing the gemcitabine IC50 levels of screened clones. Slices represent the clone selected for whole genome sequencing. Color scale is the same as panel B.

We evolved three independent populations of each *E. coli* strain in sub-inhibitory concentrations of gemcitabine (methods) and three control population without any drug. We monitored drug resistance in all population daily throughout the experiment (Figure 3B). We observed that resistance emerged within a day or two for the Nissle 1917 and F-18 strains, while it emerged more slowly for the BW25113 strain. Last day populations from all strains that evolved in gemcitabine were resistant to the drug across all tested drug concentrations (Figure 3C). In order to isolate individual resistant clones, we streaked each of the evolved population on agar plates and measured IC50 does for 8 independent colonies. (Figure 3D). Almost all clones from Nissle 1917 and F-18 drug evolved strains were resistant. However, clones from the BW25113 populations showed heterologous levels of resistance. All clones isolated from the populations that evolved without the drug were drug sensitive. We chose a single clone from each population for further analysis (marked as extruding slices in the pie charts on Figure 3D). These individual clones represent lineages that evolved completely independently from one another.

### Inactivation of *nupC* underlies evolved drug resistance

Phenotypic measurements revealed that all drug-evolved populations became highly resistant. We next sequenced the genomes of evolved clones to identify the underlying adaptive mutations and identified mutations with the BreSeq software^42^. Figure 4A shows the mutations we identified in the single clones as circa plots (concentric circles representing the bacterial chromosomes). Annotation of mutations in F-18 required careful manual inspection since its reference genome consists of 113 contigs. As the figure shows, we observed that the genomes of all the drug-evolved clones harbored *nupC* mutations while none of the control-evolved clones had such mutations. Supplementary table 4 provides the BreSeq information the mutations we identified. Importantly, while we found additional mutations beyond those in the *nupC* gene, we did not observe another gene that was repeatedly mutated across all independent replicates in all of the *E. coli* strains. The only other repeated mutation we identified involved the *yahF* gene in two lines of F-18 that also shared an identical *nupC* mutation (transposon). Inspecting the BreSeq report and the contig files led us to believe both lines shared single new junction that impacted both the *nupC* and *yahF* loci.

**Figure 4:**
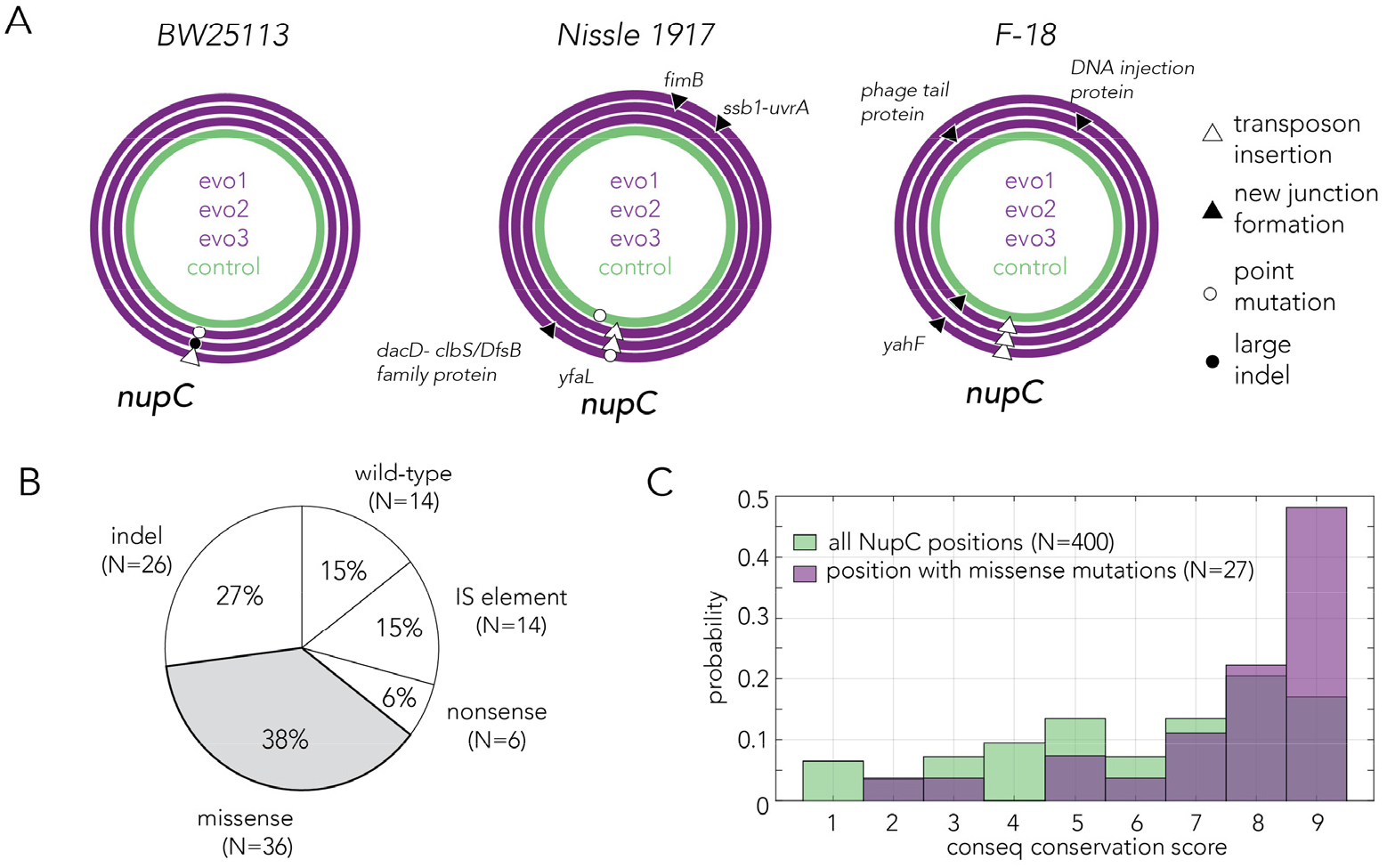
Evolved resistance converges to inactivation of the nucleoside permease NupC. **A**. Circa plots showing the mutations identified by whole-genome sequencing in pure clones isolated from independently evolved populations. Mutations in the coding region of the *nupC* gene were observed across all gemcitabine-evolved clones but not in the no-drug control evolved clones. Since the F-18 genome is not fully assembled, relative positions on the F-18 circa plot are not real genomic positions. **B**. Pie chart showing the frequency of various mutation types identified by sequencing the *nupC* across 96 spontaneous gemcitabine resistant mutants in the BW25113 strain. **C**. A comparison of the evolutionary conservation of missense mutation positions relative to the conservation of all position in NupC. The positions of the missense mutations are statistically biased towards the conserved positions (*p*-value = 1.95*10^−4^ in Wilcoxon rank sum test).

The mechanism of mutation in the *nupC* gene varied across the gemcitabine-evolved strains, and included point mutations, large deletions, and new junction formation within *nupC* coding region that can stem from transposon insertions and genomic rearrangements. The two of the point mutations we identified, were missense mutations (S175P and V249A). To evaluate if these point mutations likely interfere with the permease function, we used the ConSurf Server that identifies evolutionarily conserved positions^43^. The analysis revealed that both positions are highly conserved and are therefore likely important for the permease function (positions marked in red in supplementary Figure 5). Lastly, we examined the function of all other genes that were mutated in evolved strains to pinpoint additional putative adaptive mutations. We identified a new genomic junction, likely originating a transposon insertion, upstream to the *uvrA* gene. The *uvrA* gene codes for A subunit in the UvrABC nuclease that is involved in nucleotide excision repair pathway^44^. The mutation impacted the annotated promoter of the gene according to regulonDB^45^. Taken together, we concluded that resistance emerged across all gemcitabine-evolved strains primarily through inactivation of the nucleoside permease NupC.

### Mechanisms underlying convergence towards *nupC* inactivation

Gemcitabine adaptation in our evolution experiments likely emerged through inactivation of *nupC* across all evolved strains. This convergence can be driven by multiple mechanisms that are not necessarily exclusive to one another. Evolutionary trajectories are influenced by multiple parameters, including the adaptation benefit (e.g., the level of resistance the mutation confers), the adaptation cost (e.g., if it reduces growth) and the likelihood that the mutation will appear spontaneously. We reasoned that quantifying these parameters for the *nupC* gene would provide insight on the forces underlying the evolutionary convergence we observed.

Our genetic screen already revealed that *nupC* is among top loss-of-function mutations conferring resistance (Figure 1E), yet it remains unclear whether *nupC* inactivation is associated with any cost to the cells. We therefore monitored the growth rate of the top five resistant knockouts identified in the genetic screens (supplementary figure 4). The experiment revealed that a *nupC* knockout grows as fast as the wild-type strain (*ybcN* knockout) for most of the growth phases and slows down only during the stationary phase. In contrast, all other top resistant knockouts were significantly slower than the wild-type strain (and *nupC*) during all growth phases. We concluded that *nupC* inactivation is unique among the resistant mutations since its associated with only a small fitness cost for bacteria when the drug is not present.

We next tested whether the *nupC* locus contains a mutation hotspot. We reasoned that a mutation hotspot will be evident if we observed that multiple independent resistant clones will share a specific site or region within the *nupC* locus or will leverage on a specific mutation mechanism. We obtained independent clones by picking 96 individual colonies of the wild-type strain. We grew them overnight and plated them on agar plates with an inhibitory gemcitabine concentration (0.5 mM). We reasoned that at this high concentration, only the most resistant and fast-growing mutants, namely *nupC* mutants, will be able to form large colonies within an overnight growth. We then picked a single resistant colony from each of the 96 agar plates and sequenced the *nupC* coding sequence locus. We found that 85% of colonies were mutated in this region, with mutations spans multiple types. Missense mutations and short indels were the most frequent mutations observed (Figure 4B). In order to annotate the missense mutations and we tested if they likely disrupt positions important for permease function. We used the ConSurf sequence analysis tool and identified evolutionarily conserved positions (Supplementary Figure 5). The analysis revealed that missense mutations we identified were significantly biased towards the highly conserved regions of the permease (Figure 4C, *p*-value = 1.95*10^−4^ in Wilcoxon rank sum test).

Lastly, we decided to test if the *nupC* is naturally poised for frequent mutations in the absence of gemcitabine. We reasoned that since our agar plating experiment showed that most resistant colonies have been *nupC* mutants, we could estimate the mutation rate (μ) of in this gene with a Luria Delbrück fluctuation test^46^. We compared this mutation rate to a reference gene (*nfsA*) that confers resistance to furazolidone^47^. We performed this experiment as four replicates using four different genetic backgrounds (the strains used for the evolution experiments and the MG1655 lab strain). Supplementary table 3 shows the calculated mutation rates for *nupC* and *nfsA* loci in all strains (normalized to a gene with 1000 basepairs). In all cases we found rates comparable to the average gene mutation rate previously inferred for *E. coli* (2.1 × 10^−7^ per gene per generation)^48^. For BW25113, the mutation rate of *nupC* was 5.7-fold higher than that of *nfsA*. For Nissle, this difference was 6.8-fold and for MG1655 it was 1.8-fold. For F-18 strain, in contrast, the mutation rate of *nfsA* was 7.9-fold higher than *nupC*. While the mutation rate was not identical in the two genes, the mutation of *nupC* was not always higher than that of *nfsA*, and their rate difference was never larger by an order of magnitude.

Taken together, the results from multiple experiments suggested that *nupC* was likely repeatedly inactivated across multiple genetic backgrounds since it confers high drug resistance without compromising growth. Experiments focusing on the *nupC* mutation revealed that gene inactivation could take place through multiple alternative mutation mechanisms and that the *nupC* locus is not characterized by a exceptionally high mutation rate (excluding the possibility of a mutational hotspot).

## Discussion

The recent discovery that bacterial infections are frequent across multiple cancer types suggests that the tumor-microbiome is an important, yet under-studied, component of the tumor microenvironment^3,4^. Here, we suggest that a key aspect of microbial biology is under-explored in current investigations of the tumor-microbiome – the ability of bacteria to rapidly evolve and adapt to extracellular changes. Within the tumor niche, successful colonization may require bacterial adaptation to the unique conditions of the tumor microenvironment, including adaptation to tumor-targeting therapeutics. A strong selective pressure from chemotherapies likely exists given that multiple antineoplastic drugs are putative antimicrobials at physiological concetrations^11^. We previously raised the hypothesis that bacterial evolved resistance to tumor-targeting chemotherapies can manifest in changes to bacterial drug metabolism and this type of adaptation can inadvertently influence chemoresistance. Our previous work used the *Caenorhabditis elegans* model system, its bacterial diet, and two fluoropyrimidine chemotherapy drugs to study this hypothesis^17^. In that model system, we estimated that almost 60% of loss-of-function mutations that confer bacterial resistance will also reduce drug toxicity in a worm host feeding on these bacteria. Here we further study this hypothesis and focus on a different chemotherapy drug and a model system that captures bacterial-drug interactions that may be at play in the tumor microenvironment, rather than in the host gut as in the previous publication.

Research of the bacterial role in pancreatic cancer revealed multiple and independent mechanisms of microbial influence on cancer in this organ^4,7,20–23^. Specifically, recent work suggested that pancreatic colonization by proteobacteria can decrease efficacy of gemcitabine through rapid bacterial drug inactiavation^21^. We used this drug-bacteria-tumor interaction to test our hypothesis by reconstituting similar interactions from individual parts that are well-understood on their own. We first mapped the gemcitabine resistome in *E. coli* and found that changes in multiple cellular processes increase resistance. Importantly, these potential adaptations are easily accessible within short time scales since they only require inactivation of a single gene. A functional assay revealed that one third of these adaptations impact bacterial drug breakdown (Figure 2B). Indeed, gemcitabine administration to co-cultures of cancer spheroids and bacteria demonstrated that two of the top loss-of-function mutations that the screen identified can considerably, and oppositely, impact drug efficacy in neighboring cancer cells (Figure 2E). While the impact of bacterial resistance on neighboring cells could have been predicted for the *nupC* knockout, it was not trivial for the *cytR* knockout. CytR is a transcription factor that represses at least 14 different operons, including nucleotide transporters and membrane proteins (*nupC*, *nupG*, *tsx*, *ycdZ*), sigma factors (*rpoH*), and metabolic enzymes in the target pathways (*ccd*, *udp*, *deoABCD*)^44^. A chemical assay revealed that *cytR* knockdown culminates in increased drug degradation and a co-culture experiment confirmed it increases chemoresistance of neighboring cancer spheroids. It is therefore likely that relief of CytR repression of both *nupC* and *cdd* leads to increased import that is counteracted by even faster gemcitabine deamination. This observation is key since it suggests that mutations conferring gemcitabine resistance in bacteria can both increase and decrease gemcitabine breakdown rate. In the context of the tumor-microbiome, bacterial adaptation to gemcitabine can therefore raise or reduce the tumor’s chemoresistance.

A key question that arises from the observation that bacterial resistance can modulate drug availability for neighboring cancer cells, is which adaptations will emerge under drug selection and whether evolution will repeatedly converge to the same resistance mechanism. Results from the serial transfer evolution experiment in three *E. coli* strains showed that drug selection repeatedly yielded adapted clones that disabled the drug permease (Figure 4A). Additional experiments suggested that selection for *nupC* inactivation, at the expense of alternative resistance mechanisms, is attributed the high resistance *nupC* inactivation confers and to the minimal impact it has on growth. Importantly, as the fluctuation experiments reveal, resistant clones can preexist drug exposure and therefore also likely preexist any treatment if the bacterial number in the tumor microbiome is sufficiently large. In such cases, tumor-microbiome adaptation can take place through ecological-like changes and take-over by a resistant bacterial clone. Such clonal expansion in bacteria is reminiscent to the process of clonal expansion and takeover of preexisting resistant cancer cells that is thought to be prevalent in cancer treatment.

Previous research established that the microbiomes, in natural body sites or within tumors, can metabolize tumor-targeting drugs by doing so, influence drug efficacy in the host. Our previous and current studies complement this premise by demonstrating that bacterial evolutionary adaptation to chemotherapies can further influence bacterial drug metabolism and therefore host drug efficacy. Intriguingly, in this work we found that bacterial influence can have opposite effects on drug breakdown and therefore can increase, or decrease, chemoresistance. Moreover, the genetic screens from both works suggest that such occurrences might not be rare, given that such a high fraction of adaptive loss-of-function mutations also alter drug breakdown rates.

While we believe this work illuminates an under-explored and potentially impactful field of research - evolutionary adaption in the tumor microbiome - we are also excited by the follow-up questions that naturally ensue our in-vitro findings. Foremost, it will be fascinating to test if tumor colonizing bacteria evolve and adapt to anticancer host treatment in animal models. Specifically, following our observations in the model gammaproteobacteria *E. coli*, and the known prevalence of gammaproteobacteria colonization in pancreatic cancer^21^, it will be intriguing to check if similar adaptations are observed in bacteria isolated from tumors that were resected from pancreatic cancer patients. Identifying convergent adaptation in tumor-microbiome of patients may prove impactful for personalizing anticancer treatment and informing the decision to complement chemotherapy treatment with antibiotics, a decision that is highly consequential for cancer patients and that should therefore be well-justified^24,26,31,32^.

## Materials and Methods

### Bacterial strains and growth conditions

Bacterial strains used in this study are shown in table 1. We used the *E. coli* barcoded knockout strain collection for the pooled genetic screen (similarly to^17,34^). All experiments measuring gemcitabine breakdown were performed with strains from the KEIO strain collection^36^. For spheroid experiments, double knockout strains were generated with P1 transduction method^49^ using *pyrD* knockout strain from the barcoded library and the desired gene knockout from the KEIO collection. The *pyrD* knockout background was used since it is a pyrimidine auxotroph that can not grow in the media used to culture the spheroids.

**Table 1:**
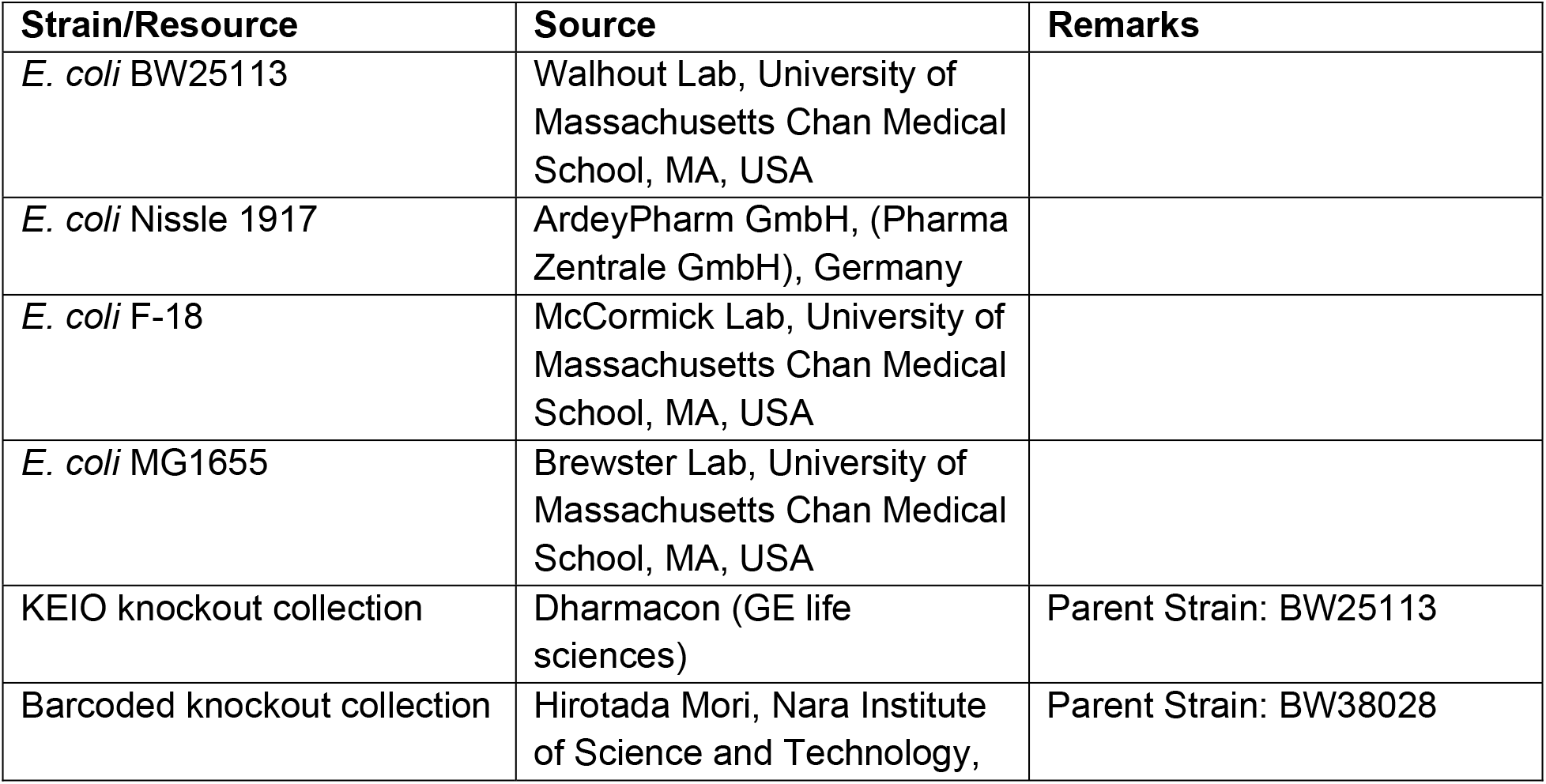
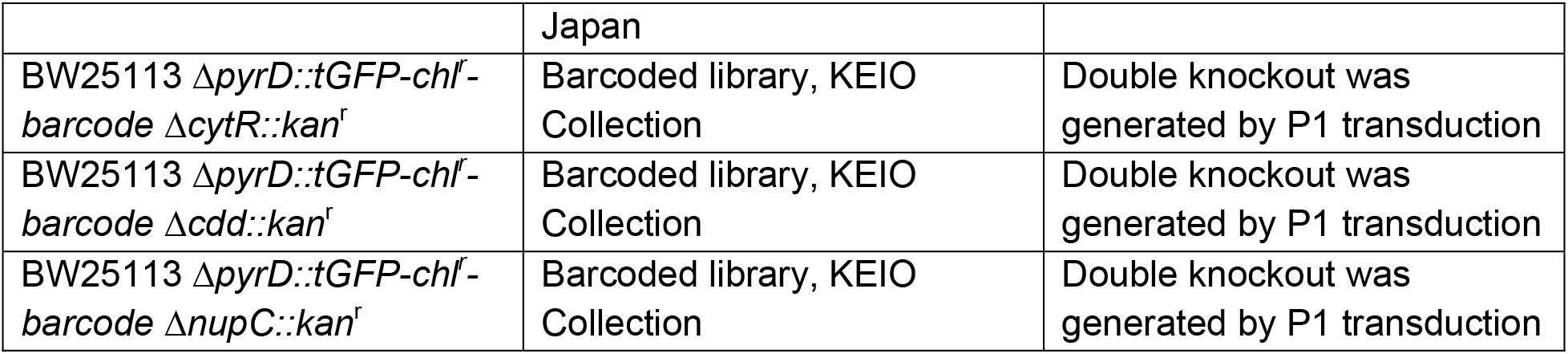
Bacteria used in this study

For all experiments, bacteria were inoculated into Lysogeny Broth (LB) and grown overnight at 37°C, 200 rpm orbital shaking. Knockout strains were grown in LB media supplemented with 50 μg/mL kanamycin (KEIO strain collection) or 25 μg/mL chloramphenicol (barcoded strain collection). All growth and serial evolution experiments were performed in M9 minimal media supplemented with 0.2% amicase and 0.4% glucose. Experiments designed to monitor gemcitabine breakdown were performed in PBS (functional assay) or in 0.9% saline (GC-MS).

### Measurement of bacterial gemcitabine dose responses and IC50

On the day of the experiment, a 384-well plate containing serially diluted gemcitabine in M9 was prepared at 2x concentration in a volume of 35 μL. For the day-to-day IC50 measurements (Figure 3B), a sample from daily evolving populations was directly diluted 1:200 into the microtiter plate with gemcitabine. For measurement of gemcitabine resistance of evolved populations (Figure 3C), a sample from frozen last day glycerol stocks of evolution experiment were inoculated into 3 mL LB for overnight culture. Overnight cultures were diluted to OD (600 nm) of 1 in M9 and were added to the 384-well plate (1:200 final dilution). For single colony gemcitabine IC50 experiments (Figure 1B and Figure 3D), single colonies were grown overnight in 3mL LB and the same dilution protocol was followed as evolved populations. The prepared microplates (with bacteria and gemcitabine dilutions) were incubated at 37°C and 360 rpm double orbital shaking in an automated plate reader (BioTek Eon) and absorbance (600 nm) was monitored every 10 minutes for 18 hours. All measurements were performed in technical triplicates. Each evolved population is considered as a separate biological replicate (three biological replicates per evolution condition. One biological replicate is shown on the figure for control evolved population.) A matlab script was used to fit the dose response curves and infer the IC50 values. Individual growth curves were assessed for quality control and to determine the exclusion criteria for analysis.

### Pooled genetic screen

We thawed a frozen glycerol stock of the pooled barcoded strain collection and inoculated 15 μL of the stock into 25 mL of LB supplemented with chloramphenicol for overnight growth at 37°C and 200 rpm shaking. In the morning, the culture was diluted to OD_600_ of 1, and then diluted 1:400 into 7ml of M9 with or without 140 μM gemcitabine. We prepared three replicates for each of the two conditions. The tubes were incubated at 37°C shaker and OD was monitored periodically. Once culture crossed OD (600 nm) 0.6, we collected the cells and extracted genomic DNA with a commercial kit (Zymo Quick DNA miniprep Plus Kit, Cat#D4068).

Library preparation was identical to the protocol we previously developed^17^. Briefly, genomic DNA isolated from endpoint of the genetic screen was quantified using Qubit dsDNA high sensitivity kit (Thermo-fisher, Cat#Q32854). We used 6.25 ng DNA to prepare the DNA library. First, barcoded region was amplified using the following primers and 2x KAPA HiFi Hotstart ReadyMix (Kapa Biosystems, Cat#KK2602) which yielded ~350 bp product. PCR products were purified using AMPure XP beads (Beckman Coulter, Cat#A63881). Nextera XT Index Kit protocol (Illumina, Cat#FC-131– 1024) was used to add indices and Illumina sequencing adapters to each PCR sample. Next, products were purified using AMPure XP bead (Beckman Coulter, Cat#A63881) purification protocol. The libraries were then run on a 3% agarose gel and the product was extracted using NEB Monarch DNA Gel Extraction Kit (NEB, Cat# T1020L). Next, we used Agilent High Sensitivity DNA Kit (Agilent Technologies, Cat# 5067–4626) to evaluate the quality and average size of the libraries. Using Qubit dsDNA high sensitivity kit, we measured the concentration and calculated the molarity of each library. Libraries were normalized to 4 nM, denatured, and diluted according to Illumina MiniSeq System Denature and Dilute Libraries Guide. After pooling, sequencing was performed using MiniSeq High Output Reagent Kit, 75-cycles (Illumina, Cat# FC-420–1001). Raw reads were deposited to NCBI Sequence read archive (SRA) under the bioproject PRJNA797841. Raw reads were converted to barcode counts using a Matlab script which compared a database of all barcodes to the reads^17^.

We identified the enriched or depleted hits by comparing the relative frequency of individual barcodes when the pooled library grew in the presence or absence of gemcitabine. For this analysis, we used the barcode counts and we identified barcodes with significant changes in their relative frequency with DEBRA^50^. We discarded barcodes with less than 10 counts. We used “Wald statistical test” and a cutoff value of 16-fold for enrichment and false discovery rate adjusted *p*-value of 0.05. Next, we performed gene set enrichment analysis with GAGE^37^ using KEGG^51^ and GO^39^ databases at a false-discovery-rate adjusted *p*-value of 0.1. We used published data on the KEIO strain collection (supplementary table 3 on ^36^) to classify slow growing knockouts. Specifically, we used the optical density measurements made after 24 hours of growth of the strain collection on minimal defined media (MOPS) and defined a cutoff value of 0.11 to discriminate normal and slow growth. We chose this cutoff value by the bimodal distribution of the density measurements in the dataset (this value separated the data into two unimodal histograms with 124 slow growing strains and 4,178 strains with normal growth).

### Rapid gemcitabine breakdown assay

We picked from the KEIO strain collection the top 88 gemcitabine resistant knockouts that were identified in the genetic screen. Knockout strains that were not found in the KEIO collection were picked from the barcoded knockout collection. The strains were grown in LB media with appropriate antibiotic at 37°C and 200 rpm shaking. We included six overnight cultures of the wild-type strain (BW25113) as controls. The next day, all strains were diluted to OD (600 nm) of 0.5 into 1 mL PBS with gemcitabine (200 μM) in a 96-deep well plate. The plate was incubated in a shaker at 37°C, 900 rpm orbital shaking. 250 μL of the supernatant was sampled after 15 and 45 minutes and filtered by spinning down at 5000G using a 96-well plate 0.22 μm filters (PALL Corporation, Cat#8119). We repeated this procedure for obtaining conditioned buffer three times on different days as independent biological replicates.

After we obtained the conditioned buffers, we evaluated the amount of residual gemcitabine in left by monitoring the growth of a gemcitabine sensitive reporter strain (*cdd* knockout). The *cdd* knockout was grown overnight in 3 mL M9 media at 37°C, 200 rpm shaking. The next day, the culture was first diluted to OD (600 nm) 1 and then further diluted 1:500 into M9 media. We aliquoted 150 μL of this culture into a 96-well plate and added 50 μL of conditioned buffer to each well. The plate was incubated at 37°C and 360 rpm double orbital shaking in an automated plate reader (BioTek Eon/TECAN). Absorbance (600 nm) was monitored every 10 minutes for 7 hours. We used the growth measurements from media supplemented with buffer after 15 minutes of incubation to identify fast degraders and growth measurements from media supplemented with buffer after 45 minutes of incubation to identify slow degraders. We used a statistical test to identify fast and slow degraders. For this test we calculated the area under the growth curve (AUC) after blank subtraction for each replicate and used a one-tailed t-test to test if the conditioned buffer from a knockout strain (three biological replicates) reduced or increased the AUC of the reporter strain compared to the buffer prepared with the wild-type strain (eighteen replicates). We used an FDR adjusted *p-value* of 0.1 as a cutoff for statistical significance.

### GC-MS measurement of gemcitabine and dFdU

We picked the knockout strains directly from frozen glycerol stock of the barcoded knockout collection and grew them overnight in 3 mL M9 media at 37°C, 200 rpm shaking. The next day, cultures were washed in saline (distilled water with 0.9% NaCl) and cultures were diluted to an OD (600 nm) of 0.125 in 1,350 μL of saline in a 96-deep well plate. Gemcitabine was added to each well to reach a final concentration of 80 μM and cultures were incubated in microplate shaker at 900 rpm and 37°C. We sampled 450 μL from the cultures at predetermined time points and filtered the samples the using 0.22 μm filters by centrifugation at 5000G for 5 minutes. We froze the conditioned supernatants at −20 °C until the GC-MS measurements were performed. This experiment was performed as three biological replicates (independent three overnight cultures and independent co-incubations).

For GC-MS measurements, first 200 μL of bacterial culture supernatants (or standard solution) were dried under vacuum. Dried samples were derivatized by adding 20 μL of pyridine and 50 μL of *N*-methyl-*N*-(trimethylsilyl) trifluoroacetamide (Sigma-Aldrich, Cat#M-132) followed by incubation for 3 hours at 37 °C. The derivatization reaction was allowed to complete for 5 hours at room temperature. Measurements were performed on an Agilent 7890B single quadrupole mass spectrometer coupled to an Agilent 5977B gas chromatograph with an HP-5MS Ultra Inert capillary column (30 m × 0.25 mm × 0.25 μm). Helium was used as carrier gas at flow rate of 1 ml/min (constant flow). The temperatures were set as follows: inlet at 230 °C, the transfer line at 280 °C, the MS source at 230 °C and quadrupole at 150 °C. 1 μL of sample was injected in a splitless mode. Initial oven temperature was set to 80 °C, held for one minute and then increased to 270 °C at a rate of 20 °C/min, then further increased to 285 at a rate of 5 °C/min. MS parameters were: 3 scans/sec with 30-500 m/z range, electron impact ionization energy 70 eV. Analytes were identified based on retention time, one quantifier and two qualifier ions that were manually selected using a reference compound. Gemcitabine was quantified as m/z 241 ion eluted at 13.14 min, 2’,2’-difluorodeoxyuridine was quantified as m/z 242 ion eluted at 11.42 min and Peak integration and quantification of peak areas were done using MassHunter software (RRID: SCR_015040).

### Spheroid experiments

We plated CT-26 mouse colon carcinoma cell-line (RRID:CVCL_7256) on 96-well low attachment plates (Costar, Cat#7007) as 4000 cells/well to form spheroids. Cells were incubated in RPMI 1640 media (Gibco, Cat#11875-093) with 2mM L-Glutamine, 5% Fetal Bovine Serum (Gibco, Cat#26140-079) and 25mM HEPES Buffer (Corning, Cat#25-060-CI). The plates were centrifuged at 3000G for 5 minutes and kept at in a tissue culture incubator with 37 °C with 5% CO_2_. After four days of spheroid growth, we serially diluted bacterial cultures into the spheroid microplate and incubated the co-culture for four hours with gemcitabine (1.6-fold serially diluted across the columns). Note that all the tested bacterial mutants in this experiment were on *pyrD* knockout background (a prymidine auxotroph) to avoid bacterial proliferation in cell culture media that does not contain any nucleotides. Next, the plate was washed with cell culture media with 50 μg/ml gentamicin three times. To achieve this, we used 96 channel handheld electronic pipette (Integra, Viaflo 96), and made use of gravity force. 100 μL media was aspirated capturing the spheroid from the bottom of the wells. After the spheroids sank to the bottom of the tips, the tips were touched to the surface of a fresh plate containing culture media with 50 μg/mL gentamicin, leaving the spheroids in the new plate. After three washes, the 96-well plate was transferred to an S3 imaging platform (Incucyte, Sartorius) which is housed inside a tissue culture incubator. The plate was imaged every six hours to track spheroid growth and validate that there was no residual contamination of resistant bacteria (evident by bacterial overgrowth). After seven days of growth, spheroids were washed once using cell culture media with 50 μg/mL gentamicin to get rid of any dead cell and cellular debris and a final microscopy image was captured. We calculated the area of individual spheroids using the Incucyte software (Segmentation sensitivity:40, Minimum Area Filter: 2000 μm^2^). A matlab script was used to make the fitness landscapes by fitting polynomial equations. The following are the steps followed: 1. Normalization by timepoint zero: we divided the last day spheroid area to day zero spheroid area (4 days post cell seeding). 2. Normalization by plate: we subtracted the minimum spheroid area from all spheroid areas and divided that value by second largest spheroid area in the plate minus minimum spheroid area 3. Fitting 3D surface and calculating EC50 lines: we fitted a mesh surface using normalized spheroid areas (2D) using a four degree polynomial function (‘poly44’). Lastly, we marked the EC50 line by calculating the coordinates of the 3D mesh surface where the values corresponded to a mid-response (value of 0.5). This experiment was performed with high resolution of conditions (12 x 8 conditions) which did not require technical replication.

### Lab evolution experiment

We evolved bacteria in sub-inhibitory doses of gemcitabine using a standard serial transfer protocol (200 μM for BW25113, 750 μM for Nissle 1917, 100 μM for F-18) in a deep 96-well plate. For each strain, a single colony was picked for each individual evolution line and grown overnight in M9 media (three biological replicates per condition). The cultures were normalized to OD (600nm) 1 and diluted 1:200 to a total volume of 1,200 μl M9 (with or without gemcitabine). The 96-well plate was incubated at 37 °C, 200 rpm shaking and was diluted 1:200 daily into fresh media for a period of 7 days (~53 generations). Resistance of evolving populations was measured daily by diluting the cultures 1:200 into a 384-well plate (35μl per well) containing serially diluted gemcitabine (prepared in M9 at 2x concentration at a volume of 35 μL). The microplate was incubated at 37°C and 360 rpm double orbital shaking in automated plate reader (BioTek Eon) and absorbance (600 nm) was monitored every 10 minutes for 18 hours. All absorbance measurements were performed in technical triplicates. All downstream experiments were performed using the frozen last day populations.

### Whole-genome sequencing and analysis of lab evolution experiment

We isolated single individual colonies from last day of the independently evolved populations by streaking them on LB agar plates. Eight colonies were selected and gemcitabine IC50 levels were determined using microtiter plate-based assay. The gDNA was extracted from selected cloned using Zymo Quick-DNA Fungal/Bacterial Miniprep Kit (Cat # 11-321). Ancestor gDNAs from the replicates of each strain were pooled at equal ratio and processed as a single sample. DNA sequencing was performed by Seqcenter (Pittsburg, PA). Seqcenter prepared libraries using Illumina DNA Prep Kit and IDT 10 basepair UDI indices. Sequencing was performed on Illumina NextSeq 2000 device (2×151 bp reads). For all samples, demultiplexing, quality control and adapter trimming was performed with bcl2fastq (Illumina) and trimgalore (Trim Galore, RRID:SCR_011847). DNA sequencing yielded a median coverage of ~120x per reference genome. We used Breseq tool to identify and annotate mutations^42^. Mixed ancestor populations were run in breseq population mode to evaluate all existing mutation variants and all other samples were run as pure clones. Following are the NCBI accession numbers for the reference genomes we used in the analysis: CP009273 for *E. coli* BW25113; CP058217.1 for *E. coli* Nissle 1917 and MLZI01000100.1 for F-18. The Breseq gdtools SUBTRACT/COMPARE was used to substract mutations that existed in the ancestor population (mutations existing 30% or more were considered) from the independently evolved clones. Then, we inspected Breseq reports to resolve unassigned junction evidence (we did not evaluate the unassigned missing coverage). Supplementary table 4 includes the mutations identified in all clones after manual inspection of the Breseq reports. Only the mutations which exist in the evolved clones but not in the ancestor were visualized on concentric circles shown in Figure 4A using circa software (OMGenomics). Since, F-18 genome consists of 113 contigs, the genomic locations shown on F-18 circa plot are undetermined. Mutations identified on contigs with low/unusual coverage were ignored (these are usually observed on small contigs which are shorter than 1 kb and frequently arise due to challenges in read mapping).

### Obtaining spontaneous nupC mutants

We cultured 96 cultures of the BW25113 strain from individual colonies overnight in LB media (96 biological replicates). In the morning 100 μL of each culture were plated with glass beads on M9 agar plates containing 0.5 mM gemcitabine. A day after, a single colony, corresponding to single spontaneous resistant mutant, was isolated from each agar plate. A 1.3 kb region spanning the entire *nupC* coding region was amplified by colony PCR and Sanger sequenced with forward (5’ TCACAGGACGTCATTATAGTG 3’) and reverse (5’ TGAGAGTAATTCATCGGCAC 3’) primers. We note that mutations in the promoter region were not sequenced or annotated by this method due to the position of the primers. However, some mutations in this region may account for some of the spontaneous resistant mutants that were not annotated as having *nupC* mutation (Figure 4B).

### Luria-Delbruck fluctuation experiment

For each strain, we inoculated 4 single colonies into 1 ml M9 medium and grew them at 37°C, 200 rpm shaking for 12 hours. Cultures were diluted to OD (600 nm) of 1 in M9. A 10^−6^ dilution of OD1 was used to determine accurate CFU/OD (600nm) by plating on LB agar plates. A 10^−4^ dilution of OD 1, was further diluted 27-fold and transferred into 10 wells of a 96-well plate as 200 μL/well (initial population size (N_0_): approximately 730 cells/well). The 96-well plate was incubated at 37°C, 1000 rpm shaking overnight. Next day, OD (600 nm) of 2 wells from the 96-well plate was measured to estimate the final population size (N_t_). Next, all cultures were diluted 1:40 into 1 mL of 0.9% saline. We plated 200 μL of each cell suspension on M9 agar plates (Plating efficiency (ɛ), 1:40) containing selective amounts of gemcitabine (0.5 mM for BW25113, Nissle 1917, MG1655; 62.5 μM for F-18) or furazolidone (1mg/mL for BW25113, Nissle 1917, MG1655; 2 mg/mL for F-18). Plates were incubated at 37°C for 18 hours and the number of colonies was determined. Then, mutation rates were calculated using the RSalvador package in R^52^. First, mutation frequency (m) was calculated using the function newton.LD.plating (and 95% confidence intervals were calculated using conf.LD.plating). Then mutation frequency (m) was divided to N_t_ to find mutation rate per generation (*p*). These numbers later normalized to a gene that is 1000 bp long.

## Materials and Data availability statement

*E. coli* barcoded library used in this study was kindly provided by Dr. Hirotada Mori. Gemcitabine/no drug evolved BW25113 and F-18 populations are available upon request. Gemcitabine/no drug evolved *E. coli* Nissle 1917 strain and its ancestors cannot be shared with third parties due to material transfer agreement conditions between University of Massachusetts Chan Medical School, USA and Ardeypharm, GmBH, Germany unless there is written permission from Ardeypharm, GmBH, Germany.

Raw sequencing reads from barcoded knockout library screen and whole genomes of evolved bacteria are available on NCBI Sequence Read Archive under the bioprojects PRJNA797841 and PRJNA855939. R and Matlab codes used in this manuscript are deposited to Github (Private Repository, request access)

## Acknowledgements

The research reported in this article was supported by NIGMS of the National Institutes of Health under award number R35GM133775 and R01AI170722 to AM, by DK068429 and R35GM122502 to AJMW. We would like to thank Dr Elizabeth Shank, Dr Hyun Youk and Dr Caryn Navarro for their comments on the manuscript. We thank Dr Hirotada Mori for providing us with the barcoded strain collection.

## Supplementary Figures and Tables

**Supplementary figure 1:**
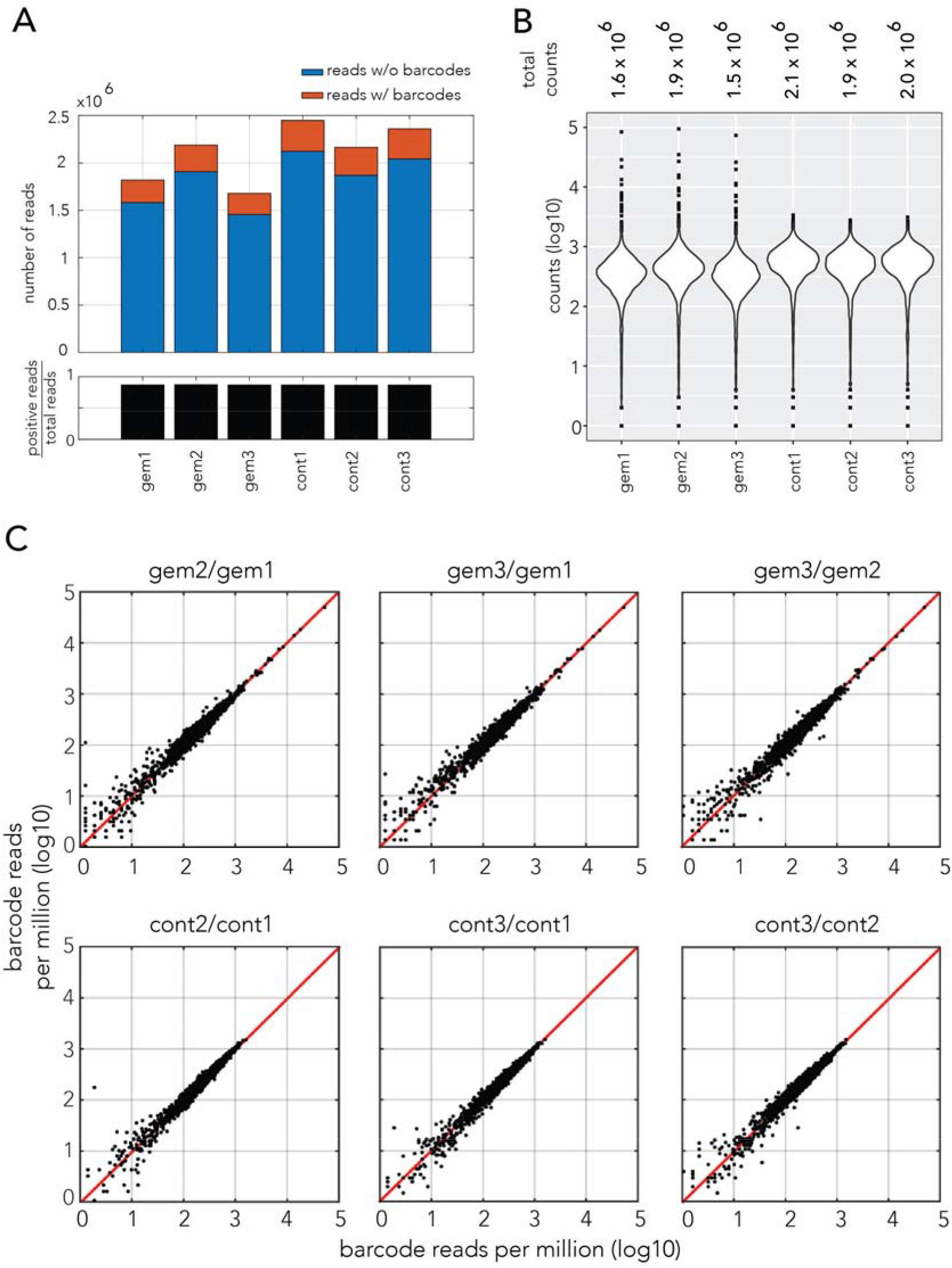
Statistics of the gemcitabine genetic screen performed with the *E. coli* barcoded knockout strain collection. **A**. Bar plots showing the number of reads with or without barcodes across samples (three biological replicates with gemcitabine and without drug). **B**. Violin plots showing the frequency of individual barcode counts. **C**. Scatter plots of barcode counts across biological replicates

**Supplementary figure 2:**
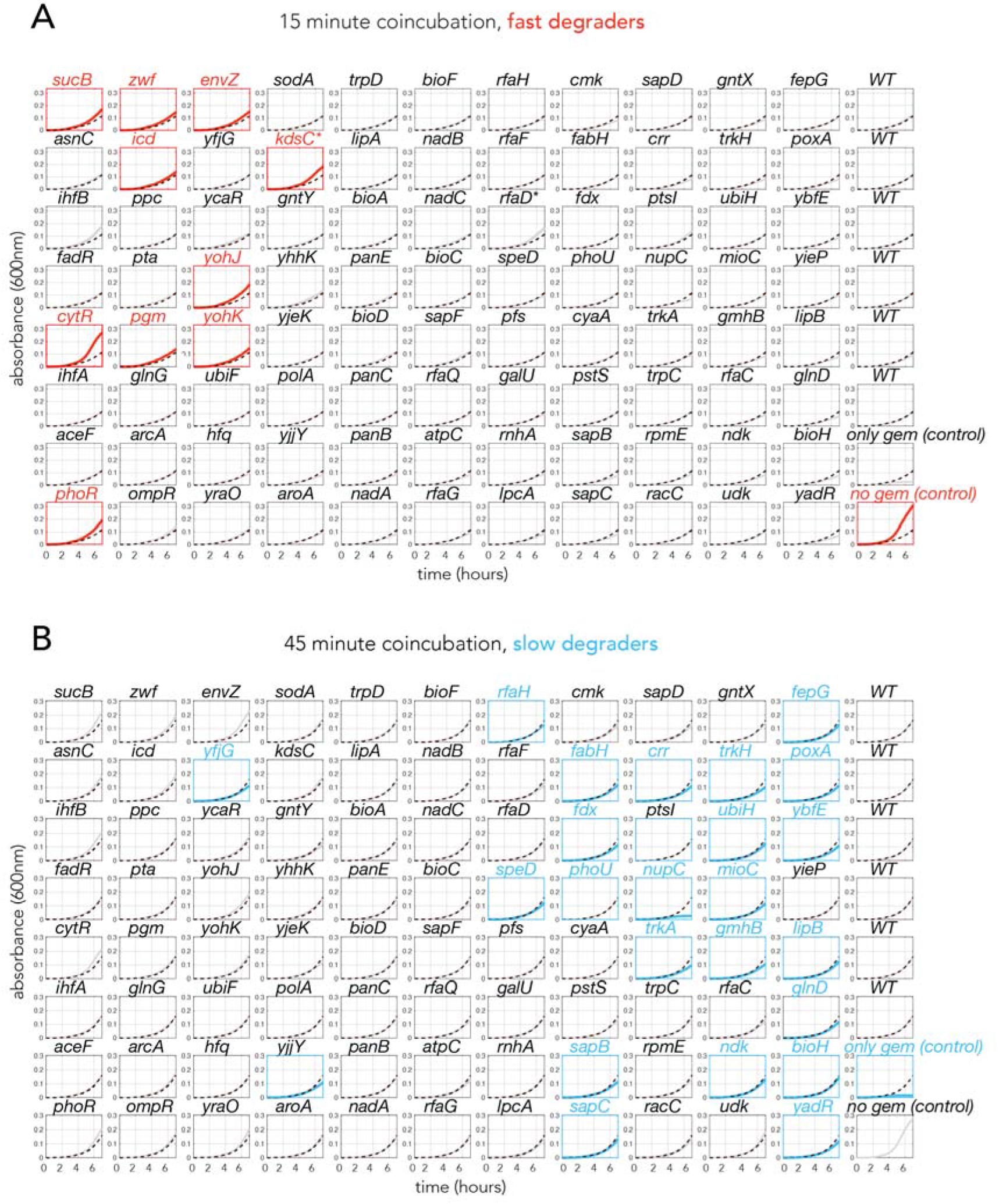
Individual growth curves of reporter strain (*cdd* knockout) in the functional assay to estimate gemcitabine breakdown rate of the top 88 gemcitabine resistant genetic screen hits. The title of each graph shows the knockout strain that was incubated with gemcitabine before the reporter strain was inoculated into the filtered conditioned supernatant. **A**. Mean growth curves of the reporter strain in the conditioned supernatant filtered after 15 min of co-incubation with the 88 tested knockout strains. Statistically significant fast degraders are shown with red color. (One tailed student’s t-test with FDR corrected *p*-adj<0.1). **B**. Mean growth curves of the reporter strain in the conditioned supernatant filtered after 45 min of co-incubation with 88 tested knockout strains. Statistically significant slow degraders are shown with blue color. (One tailed student’s t-test with FDR corrected *p*-adj<0.1).

**Supplementary figure 3:**
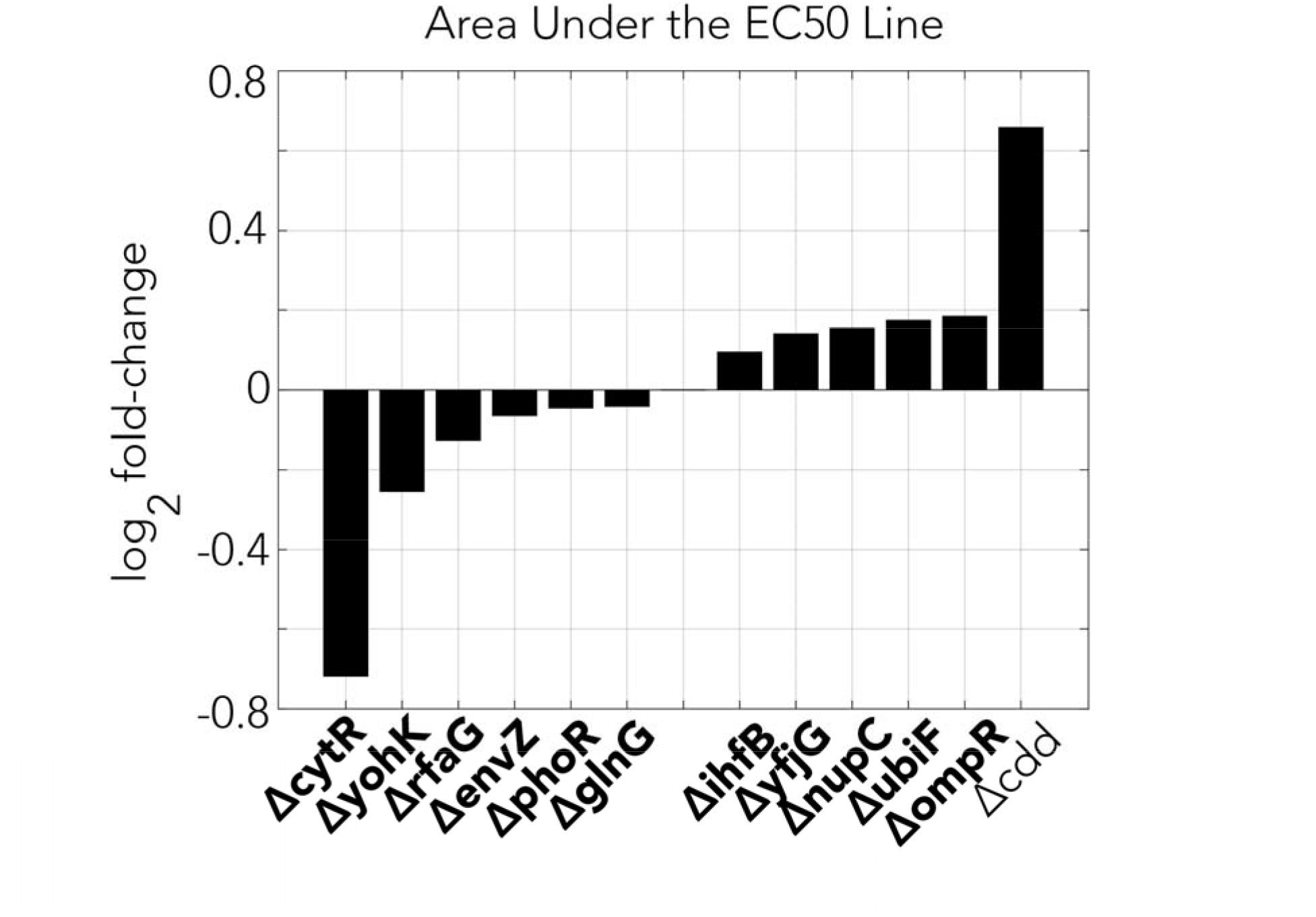
Changes in area under the EC50 front in spheroid fitness landscapes generated with co-incubation of gemcitabine and selected resistant knockout strains.

**Supplementary figure 4:**
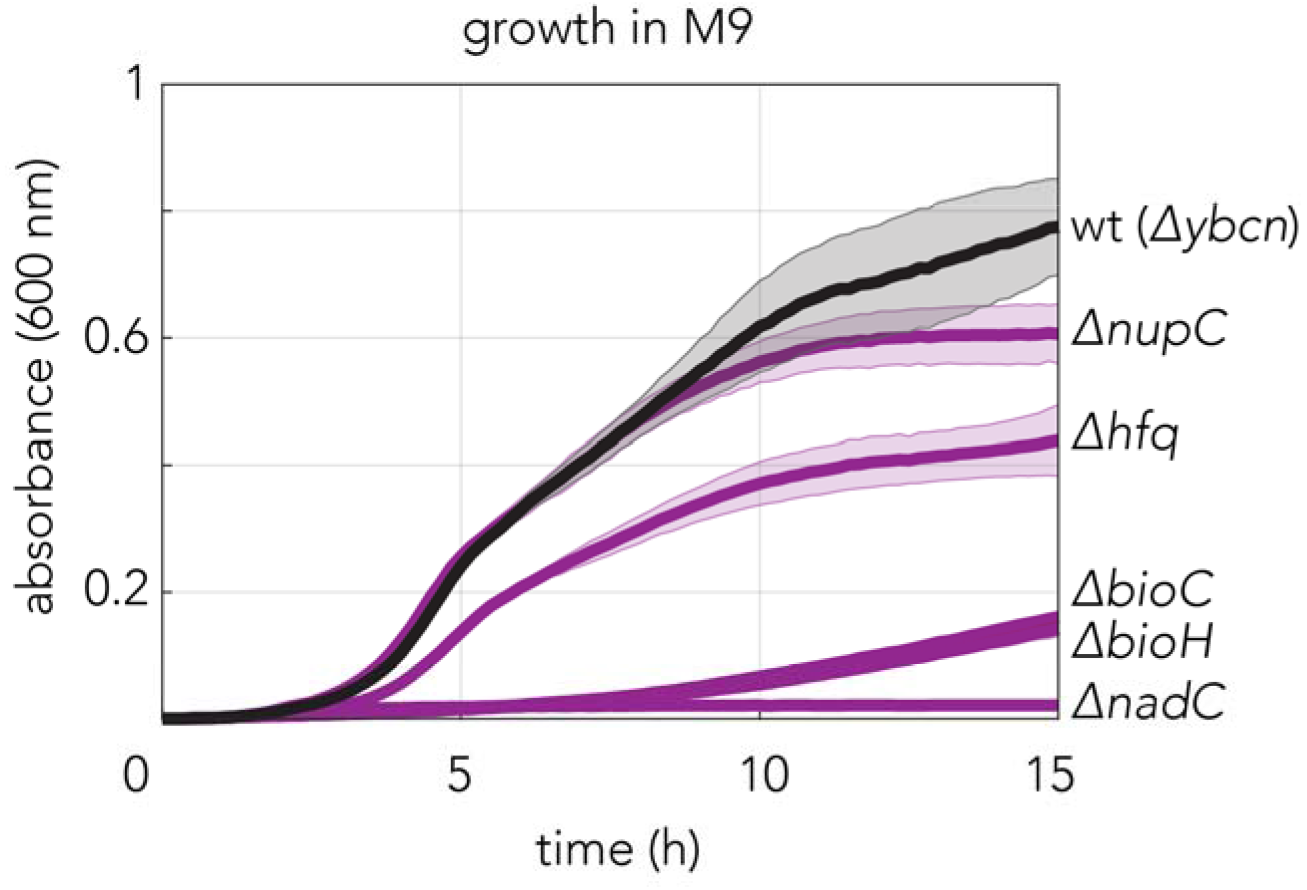
Growth of top five resistant gemcitabine knockouts indentified by the genetic screen in M9 minimal medium. The *ybcn* deletion strain was used as a wild-type control since this gene deletion does not impact *E. coli* growth and genetic background of this strain is comparable to the other mutants used in the experiment.

**Supplementary figure 5:**
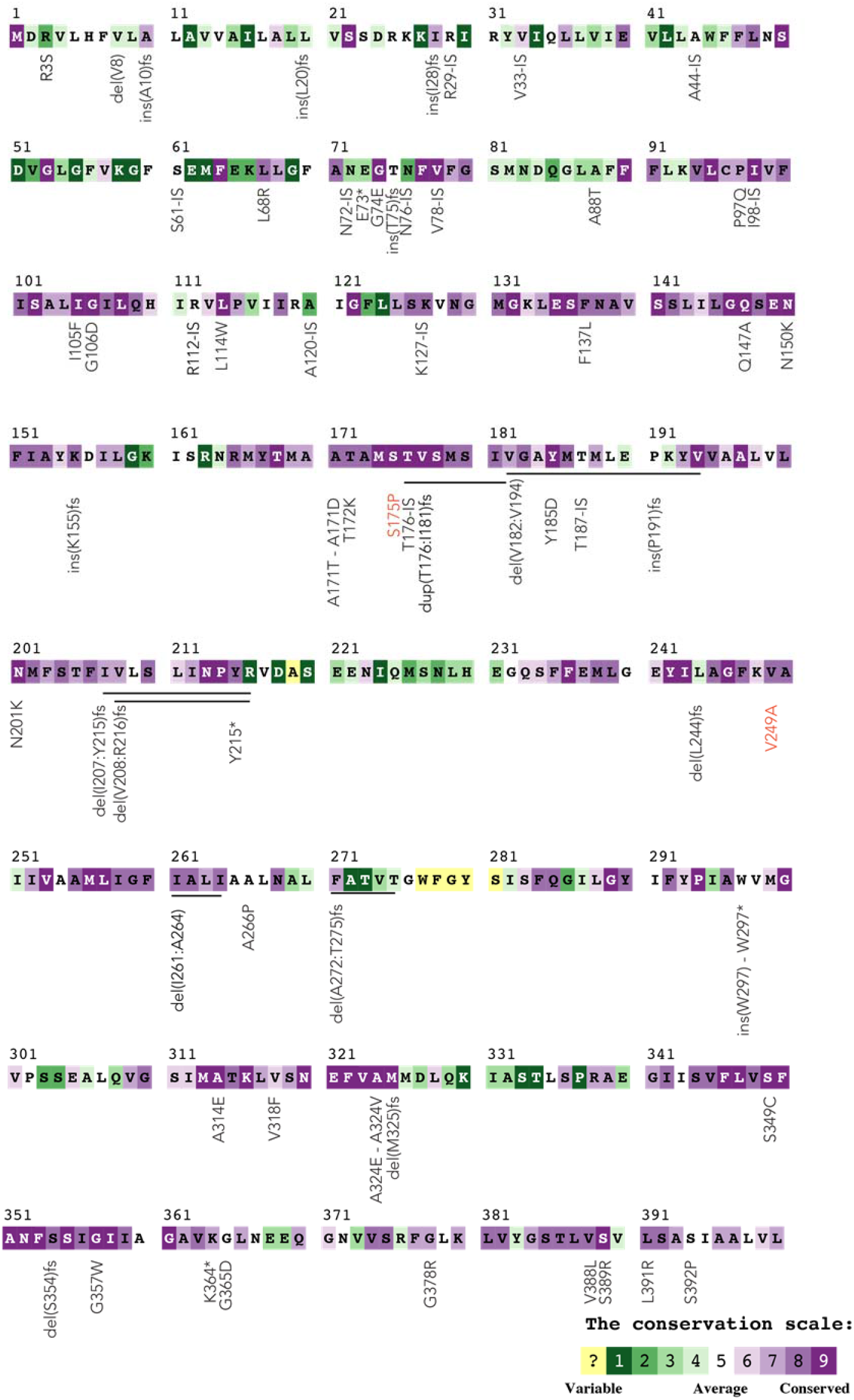
Annotation of *nupC* mutations by evolutionary conservation. The color code shows the conservation level inferred for the NupC protein with the ConSurf server. Mutations shown in black are found in spontaneous mutants isolated from agar plates containing high concentrations of gemcitabine. Mutations shown in red are missense mutations found in the gemcitabine evolved strains. Abbreviations; del:deletion, ins: insertion, fs:frameshift causing mutation, IS: insertion element(transposon insertion)

**Supplementary figure 6:**
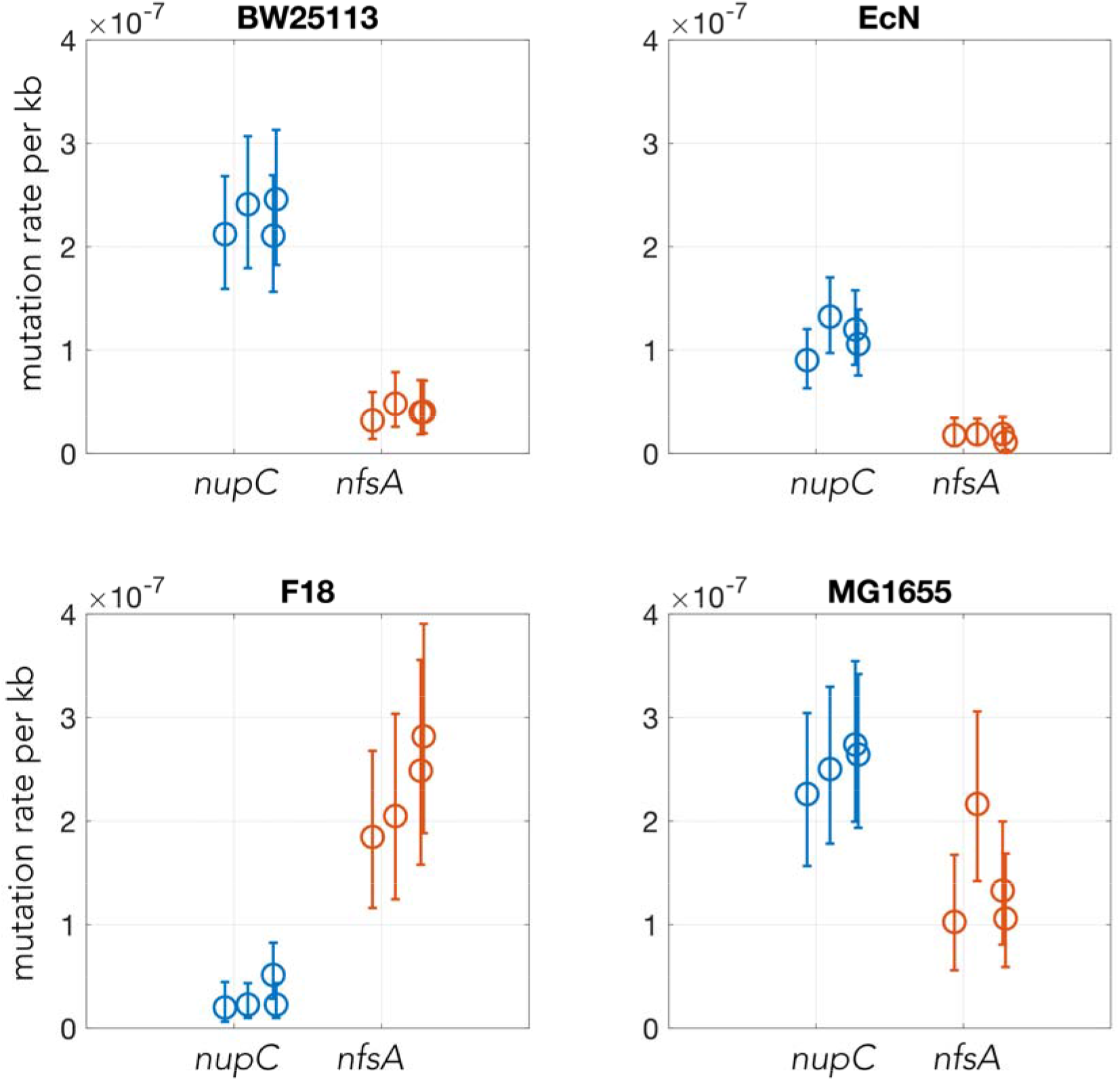
Plots showing the mutation rates in *nupC* (mutated in gemcitabine resistant mutants) and in *nfsA* (mutated in furazolidone resistant mutants) loci. The rates were calculated with a Luria-Delbrück fluctuation experiment in four *E. coli* strains. We repeated each fluctuation experiment four times.

**Supplementary table 1**: Genetic screen results. The fold-changes in strain frequency and the statistical significance were determined with DEseq2 tool.

**Supplementary table 2**: Enriched and depleted pathways in the genetic screen using two databases: KEGG:yellow, GO:green, p-adj<0.1

**Supplementary table 3**: Mutations rates for the nupC and nfsA genes determined by Luria-Delbrück fluctuation experiments. The mutation rates per locus were converted to “mutation rate per kb” to compare the two unequal size loci.

**Supplementary table 4**: Summary of the mutation detected with the BreSeq tool in evolved strains.

## References

1. Sepich-Poore, G. D. et al. The microbiome and human cancer. Science 371, eabc4552 (2021).

2. Goodman, B. & Gardner, H. The microbiome and cancer: The microbiome and cancer. J Pathology 244, 667–676 (2018).

3. Cullin, N., Antunes, C. A., Straussman, R., Stein-Thoeringer, C. K. & Elinav, E. Microbiome and cancer. Cancer Cell (2021) doi:10.1016/j.ccell.2021.08.006.

4. Nejman, D. et al. The human tumor microbiome is composed of tumor type–specific intracellular bacteria. Science 368, 973–980 (2020).

5. Roy, S. & Trinchieri, G. Microbiota: a key orchestrator of cancer therapy. Nat Rev Cancer 17, 271–285 (2017).

6. Alexander, J. L. et al. Gut microbiota modulation of chemotherapy efficacy and toxicity. Nat Rev Gastroenterology Hepatology 14, 356–365 (2017).

7. Riquelme, E. et al. Tumor Microbiome Diversity and Composition Influence Pancreatic Cancer Outcomes. Cell 178, 795–806.e12 (2019).

8. Spanogiannopoulos, P., Bess, E. N., Carmody, R. N. & Turnbaugh, P. J. The microbial pharmacists within us: a metagenomic view of xenobiotic metabolism. Nat Rev Microbiol 14, 273–287 (2016).

9. Zimmermann, M., Zimmermann-Kogadeeva, M., Wegmann, R. & Goodman, A. L. Mapping human microbiome drug metabolism by gut bacteria and their genes. Nature 570, 462–467 (2019).

10. Zimmermann, M., Patil, K. R., Typas, A. & Maier, L. Towards a mechanistic understanding of reciprocal drug–microbiome interactions. Mol Syst Biol 17, e10116 (2021).

11. Maier, L. et al. Extensive impact of non-antibiotic drugs on human gut bacteria. Nature 51, 439 (2018).

12. Zhao, S. et al. Adaptive Evolution within Gut Microbiomes of Healthy People. Cell Host Microbe 25, 656–667.e8 (2019).

13. Lieberman, T. D. Detecting bacterial adaptation within individual microbiomes. Philosophical Transactions Royal Soc B 377, 20210243 (2022).

14. Garud, N. R., Good, B. H., Hallatschek, O. & Pollard, K. S. Evolutionary dynamics of bacteria in the gut microbiome within and across hosts. Plos Biol 17, e3000102 (2019).

15. Snitkin, E. S. et al. Genomic insights into the fate of colistin resistance and Acinetobacter baumannii during patient treatment. Genome Res 23, 1155–1162 (2013).

16. Gatt, Y. E. & Margalit, H. Common adaptive strategies underlie within-host evolution of bacterial pathogens. Mol Biol Evol 38, msaa278– (2020).

17. Rosener, B. et al. Evolved bacterial resistance against fluoropyrimidines can lower chemotherapy impact in the Caenorhabditis elegans host. Elife 9, e59831 (2020).

18. Kyono, Y. et al. The Atypical Antipsychotic Quetiapine Promotes Multiple Antibiotic Resistance in Escherichia coli. J Bacteriol 204, e00102–22 (2022).

19. Alekshun, M. N. & Levy, S. B. Molecular Mechanisms of Antibacterial Multidrug Resistance. Cell 128, 1037–1050 (2007).

20. McAllister, F., Khan, M. A. W., Helmink, B. & Wargo, J. A. The Tumor Microbiome in Pancreatic Cancer: Bacteria and Beyond. Cancer Cell 36, 577–579 (2019).

21. Geller, L. T. et al. Potential role of intratumor bacteria in mediating tumor resistance to the chemotherapeutic drug gemcitabine. Science 357, 1156–1160 (2017).

22. Aykut, B. et al. The fungal mycobiome promotes pancreatic oncogenesis via activation of MBL. Nature 1–4 (2019) doi:10.1038/s41586-019-1608-2.

23. Pushalkar, S. et al. The Pancreatic Cancer Microbiome Promotes Oncogenesis by Induction of Innate and Adaptive Immune Suppression. Cancer Discov 8, 403–416 (2018).

24. Meriggi, F. & Zaniboni, A. Antibiotics and steroids, the double enemies of anticancer immunotherapy: a review of the literature. Cancer Immunol Immunother 70, 1511–1517 (2021).

25. Mohindroo, C. et al. Antibiotic use influences outcomes in advanced pancreatic adenocarcinoma patients. Cancer Med-us 10, 5041–5050 (2021).

26. Gao, Y. et al. Antibiotics for cancer treatment: A double-edged sword. J Cancer 11, 5135–5149 (2020).

27. Geller, L. T. & Straussman, R. Intratumoral bacteria may elicit chemoresistance by metabolizing anticancer agents. Mol Cell Oncol 5, 00–00 (2017).

28. Cavalcante, L. de S. & Monteiro, G. Gemcitabine: metabolism and molecular mechanisms of action, sensitivity and chemoresistance in pancreatic cancer. Eur J Pharmacol 741, 8–16 (2014).

29. Voorde, J. V. et al. Nucleoside-catabolizing Enzymes in Mycoplasma-infected Tumor Cell Cultures Compromise the Cytostatic Activity of the Anticancer Drug Gemcitabine. J Biol Chem 289, 13054–13065 (2014).

30. Lehouritis, P. et al. Local bacteria affect the efficacy of chemotherapeutic drugs. Sci Rep-uk 5, 14554 (2015).

31. Corty, R. W. et al. Antibacterial Use Is Associated with an Increased Risk of Hematologic and Gastrointestinal Adverse Events in Patients Treated with Gemcitabine for Stage IV Pancreatic Cancer. Oncol 25, 579–584 (2020).

32. Elkrief, A. et al. Antibiotics are associated with decreased progression-free survival of advanced melanoma patients treated with immune checkpoint inhibitors. Oncoimmunology 8, e1568812 (2019).

33. Yang, W. et al. Genomics of Drug Sensitivity in Cancer (GDSC): a resource for therapeutic biomarker discovery in cancer cells. Nucleic Acids Res 41, D955–61 (2012).

34. Guillen, M. N., Rosener, B., Sayin, S. & Mitchell, A. Assembling stable syntrophic Escherichia coli communities by comprehensively identifying beneficiaries of secreted goods. Cell Syst (2021) doi:10.1016/j.cels.2021.08.002.

35. Mizuno, T. & Mizushima, S. Isolation and Characterization of Deletion Mutants of ompR and envZ, Regulatory Genes for Expression of the Outer Membrane Proteins OmpC and OmpF in Escherichia coli1. J Biochem 101, 387–396 (1987).

36. Baba, T. et al. Construction of Escherichia coli K-12 in-frame, single-gene knockout mutants: the Keio collection. Mol Syst Biol 2, 2006.0008 (2006).

37. Luo, W., Friedman, M. S., Shedden, K., Hankenson, K. D. & Woolf, P. J. GAGE: generally applicable gene set enrichment for pathway analysis. Bmc Bioinformatics 10, 161 (2009).

38. Kanehisa, M. & Goto, S. KEGG: Kyoto Encyclopedia of Genes and Genomes. Nucleic Acids Res 28, 27–30 (2000).

39. Ashburner, M. et al. Gene Ontology: tool for the unification of biology. Nat Genet 25, 25–29 (2000).

40. Klünemann, M. et al. Bioaccumulation of therapeutic drugs by human gut bacteria. Nature 1–6 (2021) doi:10.1038/s41586-021-03891-8.

41. Dragosits, M. & Mattanovich, D. Adaptive laboratory evolution -- principles and applications for biotechnology. Microb Cell Fact 12, 64 (2013).

42. Barrick, J. E. et al. Identifying structural variation in haploid microbial genomes from short-read resequencing data using breseq. Bmc Genomics 15, 1039 (2014).

43. Ashkenazy, H. et al. ConSurf 2016: an improved methodology to estimate and visualize evolutionary conservation in macromolecules. Nucleic Acids Res 44, W344–W350 (2016).

44. Keseler, I. M. et al. The EcoCyc database: reflecting new knowledge about Escherichia coli K-12. Nucleic Acids Res 45, D543–D550 (2017).

45. Gama-Castro, S. et al. RegulonDB version 9.0: high-level integration of gene regulation, coexpression, motif clustering and beyond. Nucleic Acids Res 44, D133–43 (2016).

46. Luria, S. E. & Delbrück, M. Mutations of Bacteria from Virus Sensitivity to Virus Resistance. Genetics 28, 491–511 (1943).

47. Lourenço, M. et al. A Mutational Hotspot and Strong Selection Contribute to the Order of Mutations Selected for during Escherichia coli Adaptation to the Gut. Plos Genet 12, e1006420 (2016).

48. Chen, X. & Zhang, J. No Gene-Specific Optimization of Mutation Rate in Escherichia coli. Mol Biol Evol 30, 1559–1562 (2013).

49. Thomason, L. C., Costantino, N. & Court, D. L. Current Protocols in Molecular Biology. Curr Protoc Mol Biology Ed Frederick M Ausubel Et Al Chapter 1, 1.17.1–1.17.8 (2007).

50. Akimov, Y., Bulanova, D., Timonen, S., Wennerberg, K. & Aittokallio, T. Improved detection of differentially represented DNA barcodes for high-throughput clonal phenomics. Mol Syst Biol 16, e9195 (2020).

51. Kanehisa, M., Sato, Y., Kawashima, M., Furumichi, M. & Tanabe, M. KEGG as a reference resource for gene and protein annotation. Nucleic Acids Res 44, D457–62 (2016).

52. Zheng, Q. rSalvador: An R Package for the Fluctuation Experiment. G3 Genes Genomes Genetics 7, 3849–3856 (2017).

